# Inositol pyrophosphates impact phosphate homeostasis via modulation of RNA 3’ processing and transcription termination

**DOI:** 10.1101/653162

**Authors:** Ana M. Sanchez, Angad Garg, Stewart Shuman, Beate Schwer

## Abstract

Fission yeast phosphate acquisition genes *pho1*, *pho84*, and *tgp1* are repressed in phosphate-rich medium by transcription of upstream lncRNAs. Here we show that phosphate homeostasis is subject to metabolite control by inositol pyrophosphates (IPPs), exerted through the 3’-processing/termination machinery and the Pol2 CTD code. Increasing IP8 (via Asp1 IPP pyrophosphatase mutation) de-represses the *PHO* regulon and leads to precocious termination of *prt* lncRNA synthesis. *pho1* de-repression by IP8 depends on cleavage-polyadenylation factor (CPF) subunits, termination factor Rhn1, and the Thr4 letter of the CTD code. *pho1* de-repression by mutation of the Ser7 CTD letter depends on IP8. Simultaneous inactivation of the Asp1 and Aps1 IPP pyrophosphatases is lethal, but this lethality is suppressed by mutations of CPF subunits Ppn1, Swd22, Ssu72, and Ctf1 and CTD mutation T4A. Failure to synthesize IP8 (via Asp1 IPP kinase mutation) results in *pho1* hyper-repression. Synthetic lethality of *asp1*Δ with Ppn1, Swd22, and Ssu72 mutations argues that IP8 plays an important role in essential 3’-processing/termination events, albeit in a manner genetically redundant to CPF. Transcriptional profiling delineates an IPP-responsive regulon composed of genes overexpressed when IP8 levels are increased. Our results establish a novel role for IPPs in cell physiology.

## INTRODUCTION

Cells respond to phosphate starvation by inducing the transcription of phosphate acquisition genes (1, 2). The phosphate regulon in the fission yeast *Schizosaccharomyces pombe* comprises three genes that specify, respectively, a cell surface acid phosphatase Pho1, an inorganic phosphate transporter Pho84, and a glycerophosphate transporter Tgp1 (3). Expression of *pho1*, *pho84*, and *tgp1* is actively repressed during growth in phosphate-rich medium by the transcription in *cis* of a long noncoding (lnc) RNA from the respective 5’ flanking genes *prt*, *prt2*, and *nc-tgp1* (4–10). It is proposed that transcription of the upstream lncRNA interferes with expression of the downstream mRNA genes by displacing the activating transcription factor Pho7 from its binding site(s) in the mRNA promoters that overlap the lncRNA transcription units (8,10,11) (Fig. 1A).

**Figure 1.**
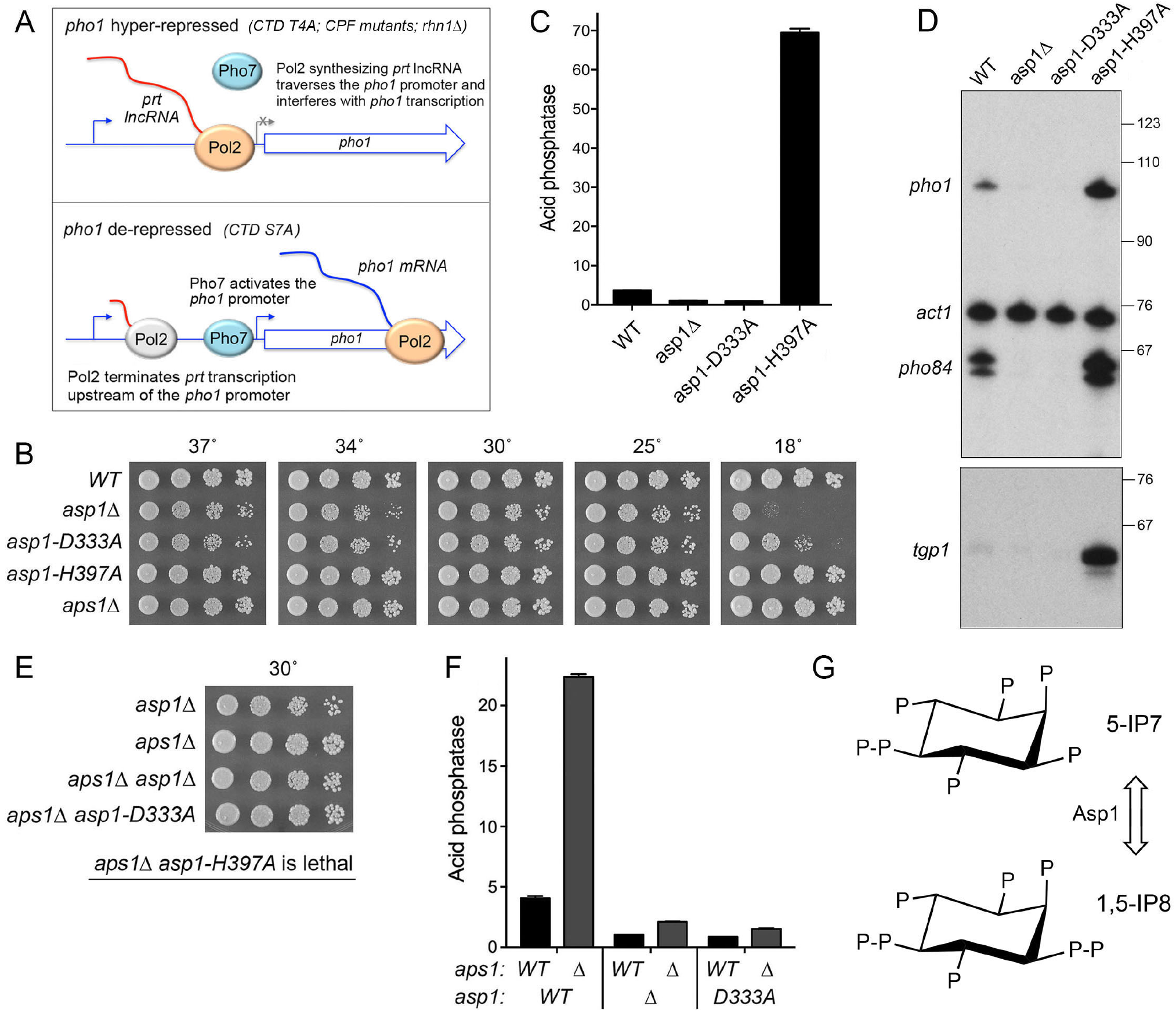
Opposite effects of IPP kinase and IPP pyrophosphatase mutations on *pho1* expression. (A) Models for the *pho1* hyper-repressed (top panel) and de-repressed (bottom panel) states of the *prt–pho1* locus under phosphate-replete conditions. (B) Growth of *S. pombe* strains with the indicated *asp1* and *aps1* alleles. Cells were inoculated in YES broth and grown at 30°C. Exponentially growing cultures were adjusted to *A*_600_ of 0.1 and aliquots (3 µl) of serial 5-fold dilutions were spotted on YES agar and then incubated at the temperatures specified. (C) The indicated fission yeast strains were grown to *A*_600_ of 0.5 to 0.8 in liquid culture in YES medium at 30°C. Cells were then harvested, washed with water, and assayed for Pho1 acid phosphatase activity by conversion of *p*-nitrophenylphosphate to *p*-nitrophenol. Activity is expressed as the ratio of *A*_410_ (*p*-nitrophenol production) to *A*_600_ (input cells). Each datum in the bar graph is the average of assays using cells from at least three independent cultures ± SEM. (D) Total RNA from fission yeast cells with the indicated *asp1* genotypes was analyzed by reverse transcription primer extension using a mixture of radiolabeled primers complementary to the *pho84*, *act1*, and *pho1* mRNAs (Top panel) or the *tgp1* mRNA (Bottom panel). The reaction products were resolved by denaturing PAGE and visualized by autoradiography. The positions and sizes (nt) of DNA markers are indicated on the right. (E) The indicated single and double mutant haploid progeny of genetic crosses were spot-tested for growth on YES agar at 30°C. The IPP pyrophosphatase-dead *aps1*Δ and *asp1-H397A* alleles were synthetically lethal. (F) The indicated strains were assayed for Pho1 acid phosphatase activity. The results show that the de-repression of Pho1 expression in *aps1*Δ cells (lacking Aps1 IPP pyrophosphatase) is erased by *asp1*Δ and *asp1-D333A* mutations that ablate Asp1 IPP kinase activity. (G) Structures of the predominant forms of IP7 and IP8 are shown. Asp1 kinase converts IP7 to IP8 and the Asp1 pyrophosphatase reverses this process (reviewed in ref. 36).

Recent studies highlight 3’ processing and transcription termination as a control point in the lncRNA-mediated repression of 3’-flanking gene expression. In particular, the *prt-pho1* locus is established as a sensitive read-out of cellular influences on termination via their effect on the degree of interference with Pho1 expression by *prt* lncRNA transcription (12). Those influences include: the phosphorylation status of the RNA polymerase II (Pol2) carboxyl-terminal domain (CTD); the activity of protein kinases Csk1 and Cdk9; the Ctf1, Ssu72, Dis2, Ppn1, and Swd22 subunits of the CPF (cleavage polyadenylation factor) complex; and the transcription termination factor Rhn1 (3,7,8,12,13).

The *S. pombe* Pol2 CTD consists of tandem repeats of a heptapeptide Y^1^S^2^P^3^T^4^S^5^P^6^S^7^. CTD mutations *S7A* and *S5A* that prevent installation of the Ser7-PO_4_ or Ser5-PO_4_ marks de-repress *pho1* and *pho84* in phosphate-replete cells (7,8,10,13). By contrast, prevention of the Thr4-PO_4_ mark by *T4A* hyper-represses *pho1* and *pho84* under phosphate-rich conditions (7, 13). Because such CTD mutations do not affect the activity of the lncRNA or mRNA gene promoters *per se* (8, 10), it is proposed that CTD status affects Pol2 termination during lncRNA synthesis (Fig. 1A). Specifically, it is hypothesized that loss of the Ser7-PO_4_ or Ser5-PO_4_ marks leads to precocious termination of *prt* lncRNA transcription prior to the *pho1* promoter and loss of the Thr4-PO_4_ mark reduces termination and hence increased transcription across the *pho1* promoter (8) (Fig. 1A).

This termination-centric scenario is supported by recent findings that: (i) mutations of CPF subunits and Rhn1 – proteins that normally promote 3’ processing/termination – result in hyper-repression of *pho1* under phosphate-replete conditions; (ii) the de-repression of *pho1* elicited by the *CTD-S7A* allele is erased by CPF and Rhn1 mutations; (iii) the *pho1* hyper-repressive *CTD-T4A* allele is synthetically lethal with the *pho1* hyper-repressive CPF alleles *ppn1*Δ and s*wd22*Δ; and (iv) the *pho1* de-repressive *CTD-S7A* allele is synthetically lethal with the *pho1* de-repressive *cdk9* alleles *T212A* and *T212E* (12). We envision that there are additional influences on Pol2 transcription termination that are amenable to discovery via their impact on fission yeast phosphate homeostasis.

Inositol pyrophosphates (IPPs) are signaling molecules implicated in a broad range of physiological processes in eukaryal cells. IPPs participate in phosphate homeostasis in *Saccharomyces cerevisiae* and *Arabidopsis thaliana* (14). A role for IPPs in *S. pombe* phosphate gene regulation was suggested by the findings that a null allele of *aps1* (which encodes a Nudix-family IPP pyrophosphatase) de-repressed Pho1 under phosphate-replete conditions, whereas a null allele of *asp1* (which encodes a kinase that synthesizes IPP) hyper-repressed Pho1 (15, 16). It was proposed that IP7 was the relevant metabolite that elicited de-repression of Pho1, based on the assumption that Asp1 kinase is responsible for IP7 synthesis from IP6 (16). This view is challenged by recent studies of the properties of fission yeast Asp1 and its effects on IPP levels *in vivo* (17–20). To wit: (i) Asp1 is actually a bifunctional enzyme composed of an N-terminal IPP-generating kinase domain and a C-terminal IPP-degrading pyrophosphatase domain; (ii) the *in vivo* effect of an *asp1*Δ null allele or a kinase-dead *asp1-D333A* allele was to eliminate intracellular IP8 and to increase the level of IP7; (iii) overexpression of the isolated Asp1 pyrophosphatase domain severely depressed intracellular IP8 and concomitantly increased IP7; and (iv) the *in vivo* effect of a pyrophosphatase-dead *asp1-H397A* allele was to increase the level of IP8 without affecting the level of IP7 (19). These results signify that the function of the Asp1 kinase is to generate IP8 via phosphorylation of its substrate IP7 and the function of the Asp1 pyrophosphatase is to convert its substrate IP8 back to IP7 (Fig. 1G).

IPP dynamics reflect a balance between a kinase that converts IP7 to IP8 and pyrophosphatases that hydrolyze IP8 to IP7. Here we focus on the how genetic perturbations of IP8/IP7 metabolism impact the expression of phosphate-responsive genes during phosphate-replete growth. Our findings implicate IP8 in promoting precocious termination of *PHO*-regulatory lncRNA transcription. Shared phenotypes, epistasis, and synthetic lethalities establish a novel interactome embracing IPPs, RNA 3’ processing and termination factors, and the Pol2 CTD.

## METHODS

### Mutational effects on fission yeast growth

Cultures of *S. pombe* strains were grown in liquid medium until *A*_600_ reached 0.6–0.8. The cultures were adjusted to a final *A*_600_ of 0.1, and 3 µl aliquots of serial 5-fold dilutions were spotted on YES agar. The plates were photographed after incubation for 2 days at 34°C, 2.5 days at 30°C and 37°C, 4 days at 25°C, 6 days at 20°C, and 8 days at 18°C.

### Deletion of asp1 and aps1

PCR amplification and standard cloning methods were employed to construct plasmids in which an antibiotic-resistance cassette (*kanMX* or *natMX*) (21, 22) is flanked by 560-to 680-bp gene-specific DNA segments corresponding to genomic sequences upstream and downstream of the *asp1* or *aps1* ORF. The disruption cassettes were excised and transfected into diploid *S. pombe* cells. Antibiotic-resistant transformants were selected and analyzed by Southern blotting to ensure correct integration of *kanMX* or *natMX* at the target locus, thereby deleting the entire *asp1* or *aps1* ORF. Heterozygous diploids were then sporulated and G418- or nourseothricin-resistant *aps1*Δ and *asp1*Δ haploids were isolated. A hygromycin-resistant *aps1*Δ strain was generated by marker switching (22).

### Tests of mutational synergies

Standard genetic methods were employed to generate haploid strains harboring mutations/deletions in two (or three) differently marked genes. In brief, pairs of haploids with null or missense mutations were mixed on malt agar to allow mating and sporulation, and the mixture was then subjected to random spore analysis. Spores (∼1,500) were plated on YES agar and also on media selective for marked mutant alleles; the plates were incubated at 30°C for up to 5 days to allow slow growing progeny to germinate and form colonies. At least 500 viable progeny were screened by replica-plating for the presence of the second (and then third) marker gene, or by sequentially replica-plating from YES to selective media. A finding that no haploids with two marker genes were recovered after 6 to 8 days of incubation at 30°C was taken to indicate synthetic lethality. [Note that by sequentially replica-plating and gauging the numbers of colonies at each step, we ensured that wild-type (unmarked) and the viable differentially marked single mutant alleles were recovered at the expected frequencies.] Growth phenotypes of viable double and triple mutant strains were assessed in parallel with the parental and wild-type cells at different temperatures (18°C to 37°C) by spotting as described above. No synergy between two mutant alleles was indicated in cases where the double-mutant grew similar to one or both of the parent strains. Synergistic growth defects were scored as *ts* if the double-mutant cells failed to grow at high temperatures at which the single mutant strains formed colonies, but grew similarly to the single mutants at lower temperatures. Double-mutant cells that exhibit a growth defect at low (but not high) temperatures relative to the parent strains were scored as *cs*.

### Acid phosphatase activity

Cells were grown at 30°C in YES medium. Aliquots of exponentially growing cultures were harvested, washed with water, and resuspended in water. To quantify acid phosphatase activity, reaction mixtures (200 µl) containing 100 mM sodium acetate (pH 4.2), 10 mM *p*-nitrophenylphosphate, and cells (ranging from 0.01 to 0.1 *A*_600_ units) were incubated for 5 min at 30°C. The reactions were quenched by addition of 1 ml of 1 M sodium carbonate, the cells were removed by centrifugation, and the absorbance of the supernatant at 410 nm was measured. Acid phosphatase activity is expressed as the ratio of *A*_410_ (*p*-nitrophenol production) to *A*_600_ (cells). The data are averages (±SEM) of at least three assays using cells from three independent cultures.

### *prt-pho1* reporter plasmids and assays

The *prt–pho1* reporter plasmid, marked with a kanamycin-resistance gene (*kanMX*), contains the tandem *prt* and *pho1* genes spanning from 1831 nucleotides upstream of the *pho1* ORF (comprising the *prt* promoter and *prt* lncRNA) to 647 nucleotide downstream of the *pho1* ORF (8). [The *prt* transcription start site is located 1198 nucleotides upstream of the *pho1* ORF, or 1147 nucleotides upstream of the *pho1* transcription start site]. For reporter activity assays and RNA isolation, *kanMX*-marked plasmids were transfected into [*prt2-pho84-prt-pho1*]Δ cells (10) and transformants were selected on YES agar medium containing 150 µg/ml G418. Single colonies of individual transformants were pooled (≥20) and grown in liquid YES+G418 medium to *A*_600_ of 0.5–0.8. Aliquots were harvested by centrifugation for acid phosphatase activity measurements as described above or for RNA analysis as described below.

### RNA analyses

Total RNA was extracted via the hot phenol method (23) from 10 to 20 *A*_600_ units of yeast cells that had been grown exponentially to *A*_600_ of 0.6 to 0.8 at 30°C. For analysis of specific transcripts by primer extension, aliquots (15 µg) of total RNA were used as templates for M-MuLV reverse transcriptase-catalyzed extension of 5’ ^32^P-labeled oligodeoxynucleotide primers complementary to the *pho84*, *pho1*, *tgp1*, or *act1* mRNAs. The primer extension reactions were performed as described previously (24) and the products were analyzed by electrophoresis of the reaction mixtures through a 22-cm 8% polyacrylamide gel containing 7 M urea in 80 mM Tris-borate, 1.2 mM EDTA. The ^32^P-labeled primer extension products were visualized by autoradiography of the dried gel. The primer sequences were as follows: *act1* 5’-GATTTCTTCTTCCATGGTCTTGTC; *pho1* 5’-GTTGGCACAAACGACGGCC; *pho84* 5’-AATGAAGTCCGAATGCGGTTGC; *tgp1* 5’-GATTCATCCCAGCCACCAGAC. For Northern blotting, aliquots (10 µg) of total RNA were resolved by electrophoresis through a 1.2% agarose/formaldehyde gel. After photography under UV light to visualize ethidium bromide-stained rRNAs and tRNAs, the gel contents were transferred to a Hybond-XL membrane (GE Healthcare) and hybridization was performed as described previously (9) using a ^32^P-labeled single-strand DNA complementary to the segment of the *prt* RNA from nucleotides +159 to +198.

### Transcriptome profiling by RNA-seq

RNA was isolated from *S. pombe* cells grown in liquid culture in YES medium at 30°C to an *A*_600_ of 0.5-0.6. Cells were harvested by centrifugation and total RNA was extracted using hot phenol. The integrity and quantity of total RNA was gauged with an Agilent Technologies 2100 Bioanalyzer. The Illumina TruSeq stranded mRNA sample preparation kit was used to purify poly(A)^+^ RNA from 500 ng of total RNA and to carry out the subsequent steps of poly(A)^+^ RNA fragmentation, strand-specific cDNA synthesis, indexing, and amplification. Indexed libraries were normalized and pooled for paired-end sequencing performed by using an Illumina HiSeq 4000 system. FASTQ files bearing paired-end reads of length 51 bases were mapped to the *S. pombe* genome (ASM294v2.28) using HISAT2-2.1.0 with default parameters (25). The resulting SAM files were converted to BAM files using Samtools (26). Count files for individual replicates were generated with HTSeq-0.10.0 (27) using exon annotations from Pombase (GFF annotations, genome-version ASM294v2; source ‘ensembl’). RPKM analysis and pairwise correlations (Figs. S4 and S6) were performed as described previously (13). Differential gene expression and fold change analysis was performed in DESeq2 (28). Cut-off for further evaluation was set for genes that were up or down by at least two-fold in mutant fission yeast strains versus wild-type, had a p-value (Benjamini-Hochberg adjusted) of ≤0.05, and had an average normalized count across all samples of ≥100. The list of genes that were dysregulated by these criteria in one or more of the mutant strains is compiled in Supplemental Table S1 (for *WT*, *asp1-H397A*, *asp1-H397A aps1*Δ *ssu72-C13S*, and *ssu72-C13S*) and Supplemental Table S2 (for *WT*, *aps1*Δ, and *asp1-D333A*).

## RESULTS

### Opposite effects of IPP kinase and IPP pyrophosphatase mutations on *pho1* expression

We surveyed fission yeast *asp1*Δ, *asp1-D333A* (IP7 kinase-dead), and *asp1-H397A* (IP8 pyrophosphatase-dead) alleles for their effects on growth on YES agar medium at 18 to 37°C (Fig. 1B) and on *pho1* expression during exponential growth at 30°C in liquid culture under phosphate-replete conditions (Fig. 1C). The *asp1-H397A* strain grew as well as the wild-type control strain at 18°C to 37°C, as gauged by colony size. By contrast, *asp1*Δ cells formed smaller colonies at 25°C to 37°C and failed to grow at 18°C. The *asp1-D333A* strain resembled *asp1*Δ, except that it was able to form small colonies at 18°C. Acid phosphatase activity (a gauge of Pho1 enzyme level that correlates with *pho1* mRNA levels, as assayed by RT-qPCR, Northern blotting, primer extension, and RNA-seq [7,8,12,13,15]) was quantified by incubating suspensions of serial dilutions of the phosphate-replete cells for 5 min with *p*-nitrophenylphosphate and assaying colorimetrically the formation of *p*-nitrophenol. The basal Pho1 activity of wild-type *rpb1-CTD* cells was hyper-repressed by 4-fold in *asp1*Δ and *asp1-D333A* cells and de-repressed by 19-fold in *asp1-H397A* cells (Fig. 1C). These results implicate: (i) increased IP8 levels (and/or increased IP8:IP7 ratio) in the *asp1-H397A* strain (19) as a trigger for de-repression of *pho1* expression; and (ii) IP8 absence and/or increased IP7 levels in the *asp1*Δ and *asp1-D333A* strains (19) as a driver of hyper-repression of *pho1* expression under phosphate-replete conditions.

By performing primer extension analysis of *pho1*, *pho84*, and *tgp1* mRNA levels in phosphate-replete wild-type, *asp1*Δ, *asp1-D333A*, *asp1-H397A* cells, we found that all three transcripts of the *PHO* regulon were increased in the IPP pyrophosphatase-defective *asp1-H397A* strain (Fig. 1D). Conversely, the *pho1*, *pho84*, and *tgp1* mRNA levels were reduced in the IPP kinase-defective *asp1*Δ and *asp1-D333A* strains (Fig. 1D). Thus, perturbations of IPP status concordantly impact the entire *PHO* axis in fission yeast.

We also queried the effect of inactivation of a different fission yeast IPP pyrophosphatase enzyme, Aps1. Aps1 is a Nudix-family hydrolase that catalyzes hydrolysis of diadenosine polyphosphates (Ap_6_A and Ap_5_A) and IPPs (IP7 and IP8) *in vitro* (29). Intracellular levels of IP7 and IP8 in fission yeast (1 to 2 µM) greatly exceed that of Ap_5_A (4 nM) (30). Studies of the effects of *aps1*Δ and *aps1* overexpression did not definitively assign a physiologic substrate, but did point to IPPs as the best candidates, rather than Ap_5_A (30). The *aps1*Δ strain grew as well as wild-type fission yeast on YES agar at 18°C to 37°C (Fig. 1B). We observed a 6-fold de-repression of Pho1 activity in *aps1*Δ cells (Fig. 1F).

### Epistasis relationships of the *asp1* and *aps1* alleles

Epistasis with respect to cell growth and *pho1* expression was queried by attempting to construct all possible combinations of double-mutant strains via mating and sporulation. We thereby found (by random spore analysis; see Methods) that *asp1*Δ *aps1*Δ and *asp1-D333A aps1*Δ strains were viable and grew well on YES agar (Fig. 1E). By contrast, the *asp1-H397A* and *aps1*Δ alleles were synthetically lethal; to wit, (i) we were unable to obtain viable double-mutants after screening a large population of haploid progeny of the genetic cross (Fig. 1E); and (ii) wild-type progeny and the differentially marked *asp1-H397A* and *aps1*Δ single mutants were recovered at the expected frequencies. This result signifies that the IPP pyrophosphatases have essential but redundant functions in fission yeast and it suggests that accumulation of too much IP8 (or too high an IP8:IP7 ratio) is in some way toxic.

Analysis of Pho1 expression in the viable double-mutants grown under phosphate-replete conditions was highly instructive, insofar as the de-repression of Pho1 elicited by the *aps1*Δ allele was erased by its combination with the hyper-repressive IPP kinase-defective *asp1*Δ and *asp1-D333A* alleles (Fig. 1F). We surmise that the de-repressive effect of loss of the Aps1 pyrophosphatase activity requires the presence of IP8 generated by the Asp1 kinase and we therefore infer that IP8 is a relevant substrate for the Aps1 pyrophosphatase with respect to phosphate homeostasis.

### Do Asp1 perturbations affect the *prt* lncRNA or *pho1* mRNA promoters?

In principle, manipulation of IPP metabolism by mutations in Asp1 could affect lncRNA-mediated repression of the downstream *PHO* genes by: (i) modulating the activity of the lncRNA promoter or (ii) affecting the intrinsic activity of the mRNA promoter (independent of lncRNA synthesis). To address these issues, we employed a plasmid-borne *prt–pho1* reporter (Fig. S1A) that was introduced into fission yeast cells in which the chromosomal *pho1* gene was deleted. This reporter faithfully reflects known homeostatic controls on the native *pho1* locus (8). Here we found that the *prt–pho1* reporter is responsive to Asp1 activity perturbations, whereby Pho1 reporter expression under phosphate-replete conditions is hyper-repressed by 4-fold in *asp1-D333A* cells and de-repressed by 8-fold in *asp1-H397A* cells compared to the wild-type *asp1*^+^ control (Fig. S1D). We proceeded to test a mutated version of the *prt–pho1* reporter construct in which the *prt* promoter is inactivated by nucleotide changes in the HomolD and TATA box elements that drive *prt* lncRNA synthesis (Fig. S1B) (8). This mutant reporter provides a readout of the intrinsic activity of the *pho1* promoter, freed from interference by transcription of the flanking *prt* lncRNA. The Pho1 activity of the mutant plasmid in wild-type cells is high (i.e., de-repressed) and is not different from the Pho1 activity in *asp1-H397A* cells (Fig. S1E). These results signify that the de-repressive effect of *asp1-H397A* on *pho1* expression from the wild-type *prt–pho1* locus is not caused by upregulation of the *pho1* promoter *per se*. Note that the Pho1 activity of the mutant reporter in *asp1-D333A* cells was 50% higher than in wild-type cells (Fig. S1E), an effect opposite to the repressive impact of *asp1-D333A* on *pho1* expression when the *prt* gene is transcribed (Fig. S1D).

The effects of Asp1 mutations on the *prt* promoter were assessed using a different plasmid reporter (Fig. S1C) in which the *prt* promoter directly drives expression of the *pho1* ORF (8). Pho1 expression from this plasmid was virtually identical in wild-type and *asp1-H397A* cells (Fig. S1F), signifying that the de-repression of native *pho1* by *asp1-H397A* is not caused by decreased activity of the *prt* promoter. The Pho1 activity from the *prt* promoter reporter in *asp1-D333A* cells was 56% higher than in wild-type cells (Fig. S1F); this effect is consonant with that seen for the *pho1* promoter reporter (Fig. S1E).

### De-repression of *pho1* expression by *asp1-H397A* and *aps1*Δ depends on CPF subunits and Rhn1

Absent an effect of the *asp1-D333A* and *asp1-H397A* alleles on the isolated *prt* and *pho1* promoters sufficient to explain their impact on phosphate homeostasis, we hypothesized that IPPs affect transcription termination during *prt* lncRNA synthesis and hence the degree of interference with the *pho1* promoter. Specifically, we envision that increased IP8 levels in the *asp1-H397A* strain lead to precocious termination during *prt* lncRNA synthesis, by enhancing the responsiveness of elongating Pol2 to the action of cleavage/polyadenylation and termination factors. If this is the case, there are clear predictions concerning epistasis relationships between *asp1-H397A* and the 3’ processing/termination machinery. Loss-of-function mutations in fission yeast proteins that promote cotranscriptional 3’ processing and transcription termination hyper-repress Pho1 under phosphate-replete conditions (12). This hyper-repression is observed in knockout strains lacking the Dis2, Ctf1, Ppn1, or Swd22 subunits of the CPF complex, a strain with a catalytically dead (C13S) version of the Ssu72 protein phosphatase subunit of CPF, and a strain that lacks the transcription termination factor Rhn1 (12; and Fig. 2A).

**Figure 2.**
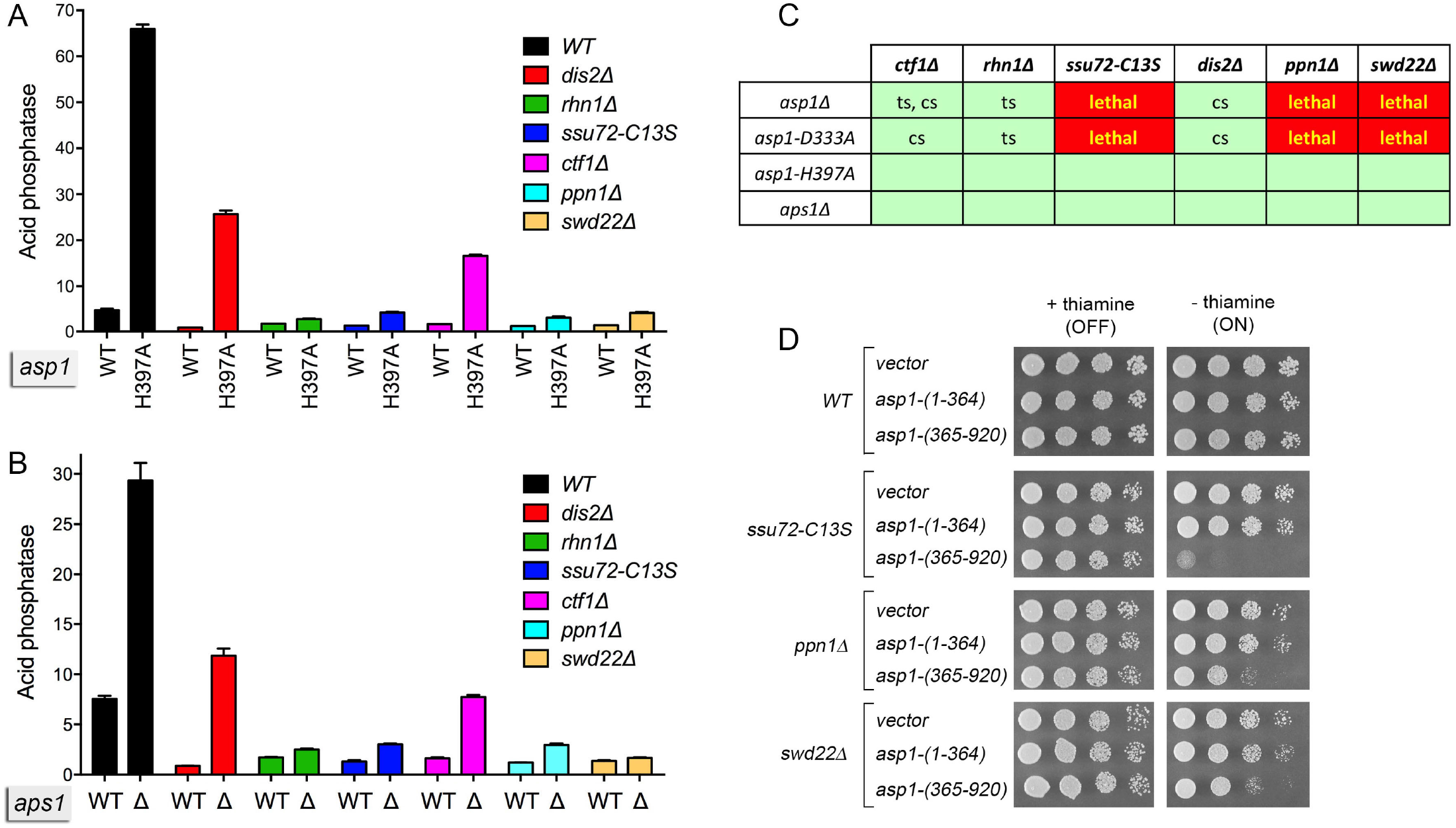
Genetic and functional interactions between Asp1, Aps1, CPF subunits and Rhn1. (A and B) De-repression of Pho1 expression by *asp1-H397A* and a*ps1*Δ depends on CPF subunits and Rhn1. *S. pombe* strains bearing the indicated *asp1* alleles (wild-type or *H397A*; panel A) or *aps1* alleles (wild-type or *aps1*Δ; panel B) in combination with CPF subunit or Rhn1 mutations as specified were grown in liquid culture at 30°C and assayed for acid phosphatase activity. (C) Synthetic lethalities and growth defects. Synthetically lethal pairs of alleles are highlighted in red boxes. Viable double-mutants without a synthetic defect are indicated by plain green boxes. Viable double-mutants that displayed temperature-sensitive (*ts*) or cold-sensitive (*cs*) defects are annotated as such in their respective green boxes. (D) Asp1 pyrophosphatase domain overexpression inhibits growth of CPF subunit mutants. Wild-type, *ssu72-C13S*, *ppn1*Δ, and *swd22*Δ strains were transformed with plasmids overexpressing the Asp1 pyrophosphatase domain (aa 365-920) or the Asp1 kinase domain (aa 1-364) under the control of the thiamine-regulated *nmt1* promoter (19). Control cells were transformed with the empty plasmid vector. After growth in PMG–Leu medium containing thiamine, serial dilutions were spotted on PMG–Leu medium that either contained 15 µM thiamine (*nmt1* promoter OFF) or lacked thiamine (*nmt1* promoter ON) and incubated at 30°C.

To address epistasis, we introduced the *asp1-H397A* allele into the CPF subunit mutant strains; the *asp1-H397A* allele did not affect their growth (Figs. 2C and 5A). It was noteworthy that introducing the *asp1-H397A* allele into the *rhn1*Δ strain suppressed the *ts* growth defect of *rhn1*Δ at 37°C (Fig. S2). We then assessed Pho1 expression under phosphate-replete conditions. The instructive findings were that the de-repression of Pho1 by *asp1-H397A* was effaced in *rhn1*Δ, *ssu72-C13S*, *ppn1*Δ, and *swd22*Δ cells and was partially blunted in *dis2*Δ and *ctf1*Δ cells (Fig. 2A). Thus, the increase in Pho1 expression in *asp1-H397A* cells requires CPF subunits and Rhn1, consistent with the precocious termination model cited above.

We extended this epistasis analysis by introducing *aps1*Δ into the various CPF subunit and Rhn1 mutant backgrounds. Note that *aps1*Δ also suppressed the *ts* growth defect of *rhn1*Δ (Fig. S2). The de-repression of Pho1 by *aps1*Δ was eliminated in *rhn1*Δ, *ssu72-C13S*, *ppn1*Δ, and *swd22*Δ cells and attenuated in *dis2*Δ and *ctf1*Δ cells (Fig. 2B). These results echo the effects of CPF and Rhn1 mutations on phosphate homeostasis in *asp1-H397A* cells (Fig. 2A) and fortify the precocious termination model.

### Effect of IPP pyrophosphatase-dead mutations on *prt* lncRNA

The *prt* lncRNA derived from the chromosomal *prt–pho1* locus in logarithmically growing vegetative cells is rapidly degraded by the nuclear exosome under the direction of DSR (determinant of selective removal) elements in the *prt* RNA (4,5,8). However, the increased gene dosage of the *prt–pho1* cassette on the reporter plasmid in *pho1*Δ cells has permitted analysis of internally terminated *prt* transcripts by Northern blotting and the identification of two internal *prt* poly(A) sites, PAS and PAS2, by 3’-RACE (12). The *prt* locus gives rise to three classes of poly(A)^+^ RNA: (i) a ∼2.5 kb RNA corresponding to a *prt–pho1* read-through transcript; (ii) a ∼0.4 kb species, *prt* PAS, that corresponds to *prt* RNA that was cleaved and polyadenylated at the +351 PAS site; and (iii) a ∼0.6 kb species, *prt* PAS2, that corresponds to *prt* RNA that was cleaved and polyadenylated at the +589 PAS2 site (Fig. 3A). These three classes of transcript are seen here in a Northern blot of RNAs isolated from two independent cultures of *asp1^+^ pho1*Δ cells bearing the *prt–pho1* reporter plasmid (Fig. 3B, lanes WT). We find that the *prt–pho1* read-through transcript is strongly suppressed in reporter-bearing *asp1-H397A* and *aps1*Δ cells, whereas the internally terminated transcripts are comparatively spared (Fig. 3B). This result is consistent with the idea that increased IPP levels in IPP pyrophosphatase-dead mutants enhances the propensity of Pol2 to terminate *prt* transcription prior to traversal of the *pho1* gene. Yet, it is not the case that the decrement in the long *prt-pho1* read-through transcript in IPP pyrophosphatase-dead mutants is accompanied by an increase in the steady-state levels of the short *prt* PAS and *prt* PAS2 RNAs. It is conceivable that the *asp1-H397A* and *aps1*Δ mutations elicit termination/polyadenylation at diffuse sites within the *prt* gene (precluding detection as discrete species on a Northern blot) or that these alleles promote turnover of transcripts precociously terminated at PAS and PAS2.

**Figure 3.**
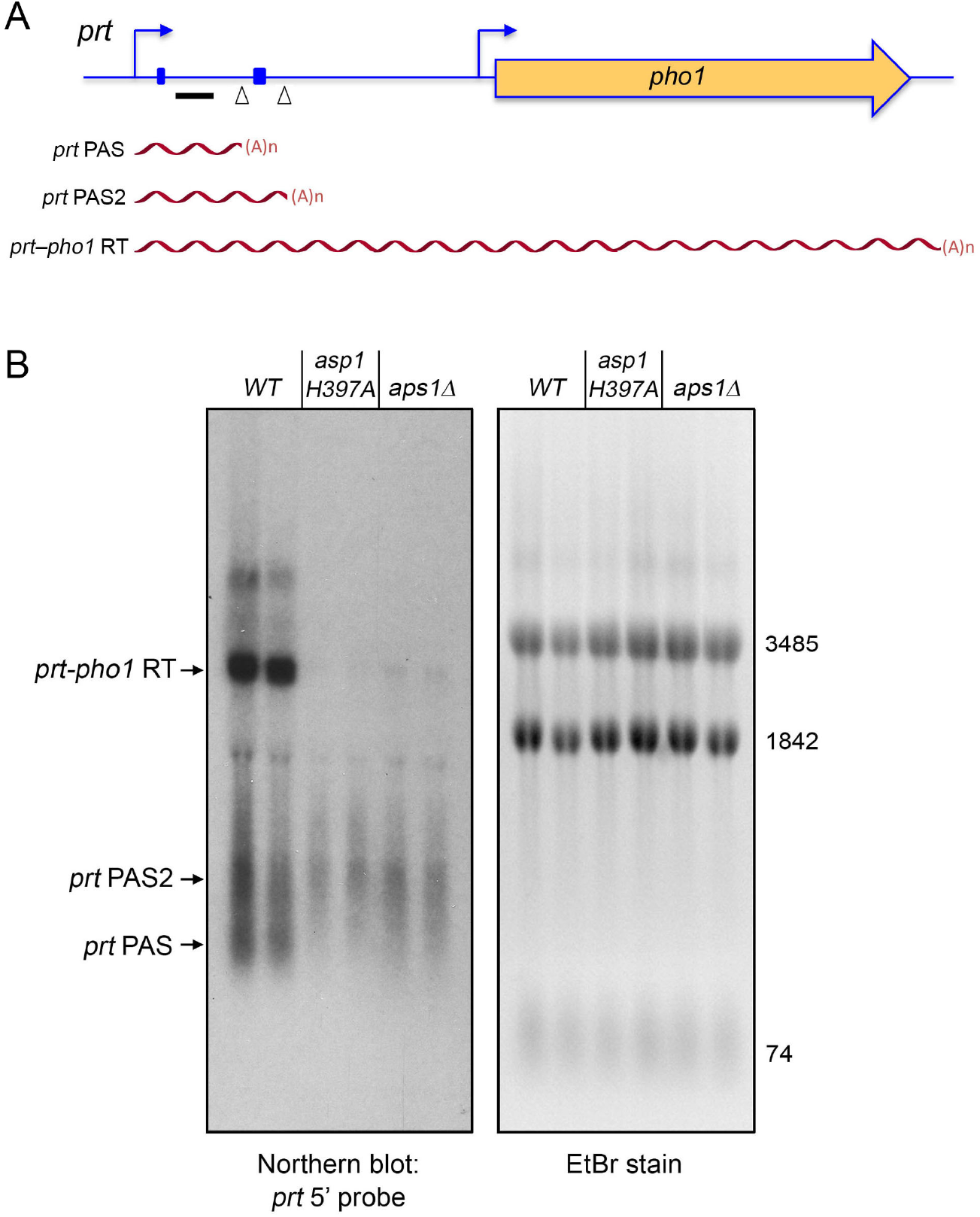
Northern analysis of *prt* RNA in IPP pyrophosphatase-dead mutants. (A) Schematic of the *prt–pho1* locus in the reporter plasmid. Transcription start sites are indicated by bent blue arrows. Triangles denote internal poly(A) sites PAS and PAS2. DSR element clusters are indicated by small blue boxes. The gene-specific probe for *prt* (a ^32^P-labeled ssDNA complementary to the segment of the *prt* RNA from nucleotides +159 to +198) is denoted by a horizontal black bar. Three classes of poly(A)^+^ *prt* transcripts described previously (12) are depicted as red wavy lines below the *prt*–*pho1* locus. (B) RNA was isolated from two independent cultures of *pho1*Δ cells bearing the *prt–pho1* reporter plasmid; the cells were either wild-type with respect to the Asp1 and Aps1 IPP pyrophosphates or were pyrophosphatase-defective *asp1-H397A* or *aps1*Δ mutants as indicated. Right panel. The RNAs were resolved by formaldehyde-agarose gel electrophoresis and stained with ethidium bromide to visualize 28S and 18S ribosomal RNAs (3485 and 1842 nucleotides, respectively) and tRNAs (74 nucleotides). Left panel. The RNAs in the gel were transferred to membrane and hybridized to the *prt* probe. Annealed probe was visualized by autoradiography. The three classes of *prt* transcripts are indicated on the left. The PAS and PAS2 transcripts are assuredly derived from the *prt–pho1* cassette, insofar as control experiments showed that they are undetectable by Northern analysis of RNA isolated from yeast strains bearing the empty plasmid vector.

### De-repression of *pho1* by *asp1-H397A* depends on DSR elements in the *prt* lncRNA

The DSR clusters in the *prt* transcript are implicated in *prt* termination (4). Therefore we tested a version of the *prt–pho1* reporter (Fig. 4A) in which the consensus DSR sequences in the two DSR clusters are altered by base substitutions (8). A *prt*-probed Northern blot of RNAs isolated from three independent cultures of *asp1^+^ pho1*Δ and *asp1-H397A pho1*Δ cells showed that formation of the *prt–pho1* read-through transcript was restored in the *asp1-H397A* strain by virtue of the DSR point mutations (Fig. 4B, compare to Fig. 3B). Moreover, the de-repression of Pho1 expression elicited by the pyrophosphatase-dead *asp1-H397A* allele was erased by the DSR point mutations (Fig. 4C). The evidence above collectively points to lncRNA termination as the target of the IPP pyrophosphatase-dead mutations on phosphate homeostasis.

**Figure 4.**
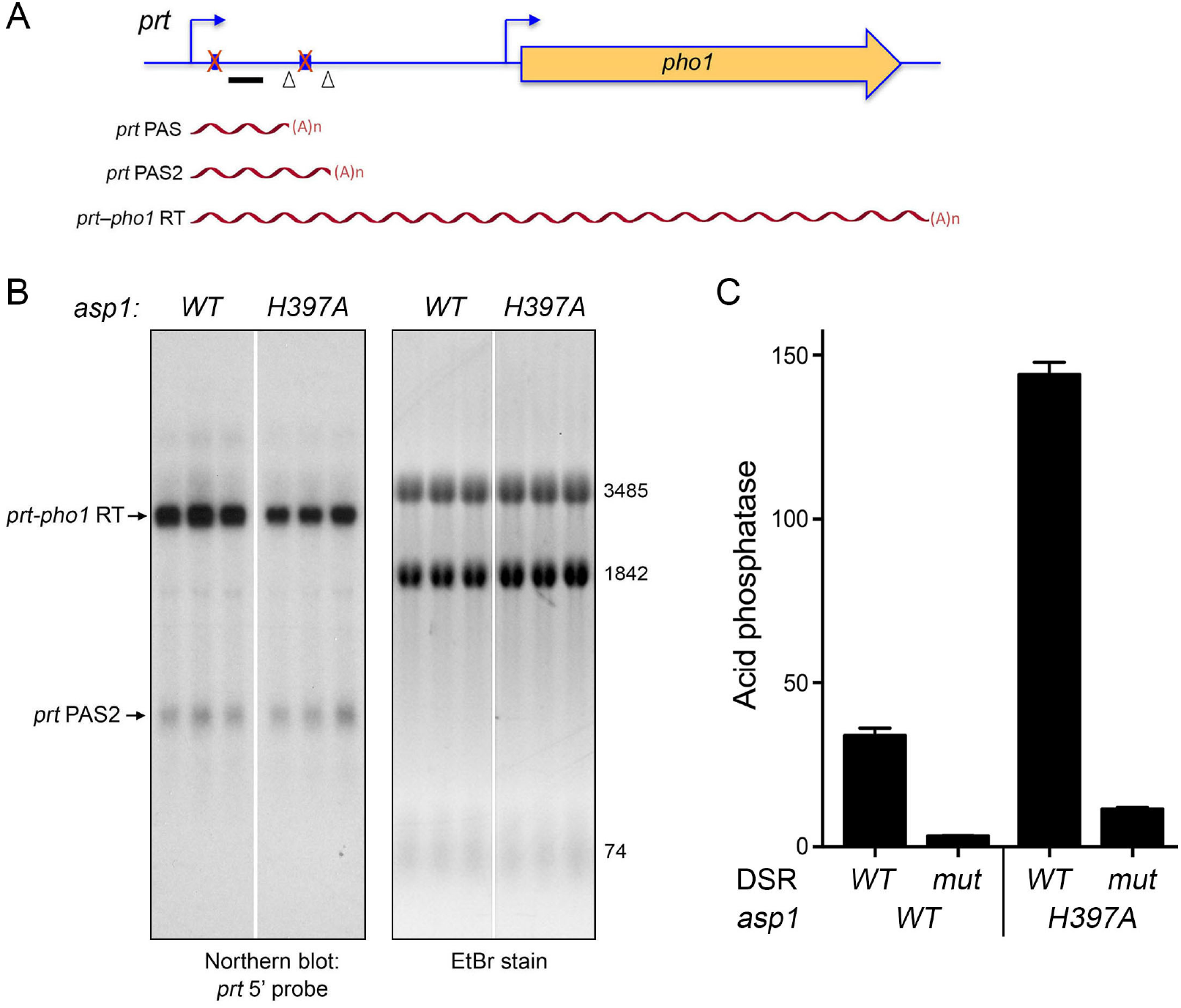
Northern analysis of *prt* RNA from a *prt–pho1* reporter with mutant DSRs. (A) Schematic of the *prt–pho1* locus in the reporter plasmid as in Fig. 3, except that both DSR clusters are mutated (indicated by a red X over the blue boxes) as described (3). (B) RNA was isolated from three independent cultures of *pho1*Δ cells (either *asp1 WT* or *H397A* as specified) bearing the DSR mutant *prt–pho1* reporter plasmid. Right panel. The RNAs were resolved by formaldehyde-agarose gel electrophoresis and stained with ethidium bromide to visualize 28S and 18S ribosomal RNAs (3485 and 1842 nucleotides, respectively) and tRNAs (74 nucleotides). Left panel. The RNAs in the gel were transferred to membrane and hybridized to the *prt* probe. Annealed probe was visualized by autoradiography. The *WT* and *asp1-H397A* RNAs were analyzed in parallel on the same agarose gel and blotted on the same membrane; the white space indicates that intervening lanes were cropped to bring the RNAs of interest close together for the purpose of composing the figure. (C) The reporter plasmids with wild-type or mutated DSRs were transfected into *asp1*^+^ (WT) or *asp1-H397A* strains in which the chromosomal *pho1* locus was deleted. Acid phosphatase activity was determined as described in Fig. S1. Each datum in the bar graph is the average of assays using cells from three independent cultures ± SEM.

### Synthetic genetic interactions of *asp1* alleles with CPF and Rhn1 mutants

To further probe genetic connections between the IPP kinase and pyrophosphatase activities and the 3’-processing/termination machinery, haploid strains with the *asp1*Δ, *asp1-D333A*, *asp1-H397A*, and *aps1*Δ alleles (marked with a 3’ flanking *natMX* or *kanMX* gene) were mated to haploid strains with null or missense mutations in cleavage/termination factors (marked with 3’ flanking antibiotic-resistance or *ura4* genes). The resulting heterozygous diploids were sporulated and, for each allelic pair, a random collection of 500-1000 viable haploid progeny were screened by serial replica-plating for the presence of the flanking markers. A failure to recover any viable haploids with both markers was taken as evidence of synthetic lethality between the test alleles. [In such cases, the wild-type haploid progeny and the differentially marked single mutants were recovered at the expected frequencies.] The lethal allelic pairs are indicated by red boxes in the matrix shown in Fig. 2C. The double-mutant haploids that passed selection were spotted on YES agar at 20°C to 37°C in parallel with the component single mutants. Viable double-mutants without a synthetic defect are indicated by plain green boxes in Fig. 2C. Viable double-mutants that displayed temperature-sensitive (*ts*) or cold-sensitive (*cs*) defects are annotated as such in their respective green boxes.

The results of the synthetic genetic array highlight allele-specific interactions of IPP metabolizing enzymes that provide new insights into the function of IPPs in fission yeast. To wit, we see that *asp1-H397A* and *aps1*Δ displayed no synthetic growth defects with the CPF subunit or Rhn1 mutants, consistent with our inferences from the phosphate homeostasis experiments that inactivation of IPP pyrophosphatases triggers precocious *prt* transcription termination and the CPF and Rhn1 mutants diminish *prt* termination. The finding that the two IPP pyrophosphatase-dead alleles reversed the *ts* growth defect of *rhn1*Δ (Fig, S2) suggests that increased IP8 ameliorates the impact of Rhn1 absence. By contrast, the IPP kinase-absent *asp1*Δ and *asp1-D333A* alleles – which we suggest are antagonists of lncRNA termination in the *prt–pho1* system – are synthetically lethal with null alleles of CPF subunits Ppn1 or Swd22 and when the Ssu72 protein phosphatase is crippled (Fig. 2C). Moreover, the *asp1*Δ and *asp1-D333A* mutations elicit conditional growth defects (*ts* or *cs*) in the *ctf1*Δ, *dis2*Δ, and *rhn1*Δ genetic backgrounds (Fig. 2C). We surmise from the synthetic lethality observed here that IP8 (synthesized by the Asp1 kinase) and the Ppn1/Swd22/Ssu72 subunits of CPF play important but genetically redundant roles in promoting essential 3’ processing/transcription termination events in fission yeast.

A prediction of the latter inference is that depletion of Asp1-synthesized IPPs via another means (not involving crippling of the Asp1 kinase) would elicit a synthetic phenotype in one or more of the CPF mutant backgrounds. Previous studies showed that this state could be achieved in *asp1*^+^ cells by overexpressing the Asp1 pyrophosphatase domain (aa 365-920) under the control of the thiamine-regulated *nmt1* promoter, which depletes intracellular IP8 and concomitantly increases IP7 (19). We found that whereas overexpression of Asp1-(365–920) under thiamine deprivation had no effect on wild-type fission yeast growth on agar medium, it prevented growth of *ssu72-C13S* cells, and slowed growth of *ppn1*Δ and *swd22*Δ cells (Fig. 2D). Controls showed that growth inhibition was specific for Asp1-(365–920), insofar as: (i) there was no effect of the *nmt1* vector or of *nmt1*-driven expression of the Asp1 kinase domain, Asp1-(1–364); and (ii) growth inhibition by the Asp1 pyrophosphatase domain was seen in the absence of thiamine when the *nmt1* promoter is on, but not in the presence of thiamine when the promoter is off (Fig. 2D).

### Lethality of *asp1-H397A aps1*Δ is rescued by *ppn1*Δ, *swd22*Δ, *ssu72-C13S*, and *ctf1*Δ

As noted above, the synthetic lethality of *asp1-H397A* and *aps1*Δ implies that accumulation of too much IP8 is in some way toxic to fission yeast. Given the potentially broad impact of IPP signaling on cellular physiology, the key question in the present context is whether the lethality of the *asp1-H397A aps1*Δ strain arises from unconstrained precocious transcription termination. If this were the case, then it might be expected that the deleterious effects of too much IP8 in the *asp1-H397A aps1*Δ context would be ameliorated by null alleles of CPF subunits Ppn1 or Swd22 or the catalytically dead CPF subunit Ssu72-C13S (each of which is synthetically lethal in the context of a kinase-dead Asp1). To test this idea, we crossed *asp1-H397A ppn1*Δ with *aps1*Δ *ppn1*^+^, *asp1-H397A swd22*Δ with *aps1*Δ *swd22*^+^, and *asp1-H397A ssu72-C13S* with *aps1Δ ssu72^+^*, then sporulated the resulting diploids, and screened random spores for each of the differentially marked loci of interest. In this way, we recovered viable *asp1-H397A aps1Δ ppn1*Δ, *asp1-H397A aps1Δ swd22*Δ, and *asp1-H397A aps1Δ ssu72-C13S* haploid strains. (By contrast, we recovered no viable *asp1-H397A aps1Δ ppn1*^+^, *asp1-H397A aps1Δ swd22*^+^, or *asp1-H397A aps1Δ ssu72*^+^ haploids.) The *asp1-H397A aps1Δ ppn1*Δ and *asp1-H397A aps1Δ swd22*Δ cells grew well on YES agar at 30°C to 37°C, but were slow growing at 25°C (Fig. 5A). The *asp1-H397A aps1Δ ssu72-C13S* strain grew poorly at 30°C to 37°C and was grossly defective at 25°C (Fig. 5A). We extended this approach to the null allele of CPF core subunit Ctf1 and recovered a viable *asp1-H397A aps1Δ ctf1*Δ haploid strain that grew slowly at 30°C and was very sick at higher and lower temperatures (Fig. 5A). These results suggest that the synthetic lethality of *asp1-H397A aps1*Δ is a consequence of IP8-driven precocious termination that depends on the Ppn1, Swd22, and Ctf1 subunits of CPF and the Ssu72 phosphatase activity. Whereas viability of *asp1-H397A aps1*Δ was restored when the aforementioned CPF subunits are deleted or inactive, additional genetic crosses showed that null alleles of Dis2 or Rhn1 were unable to suppress the synthetic lethality of *asp1-H397A aps1*Δ.

**Figure 5.**
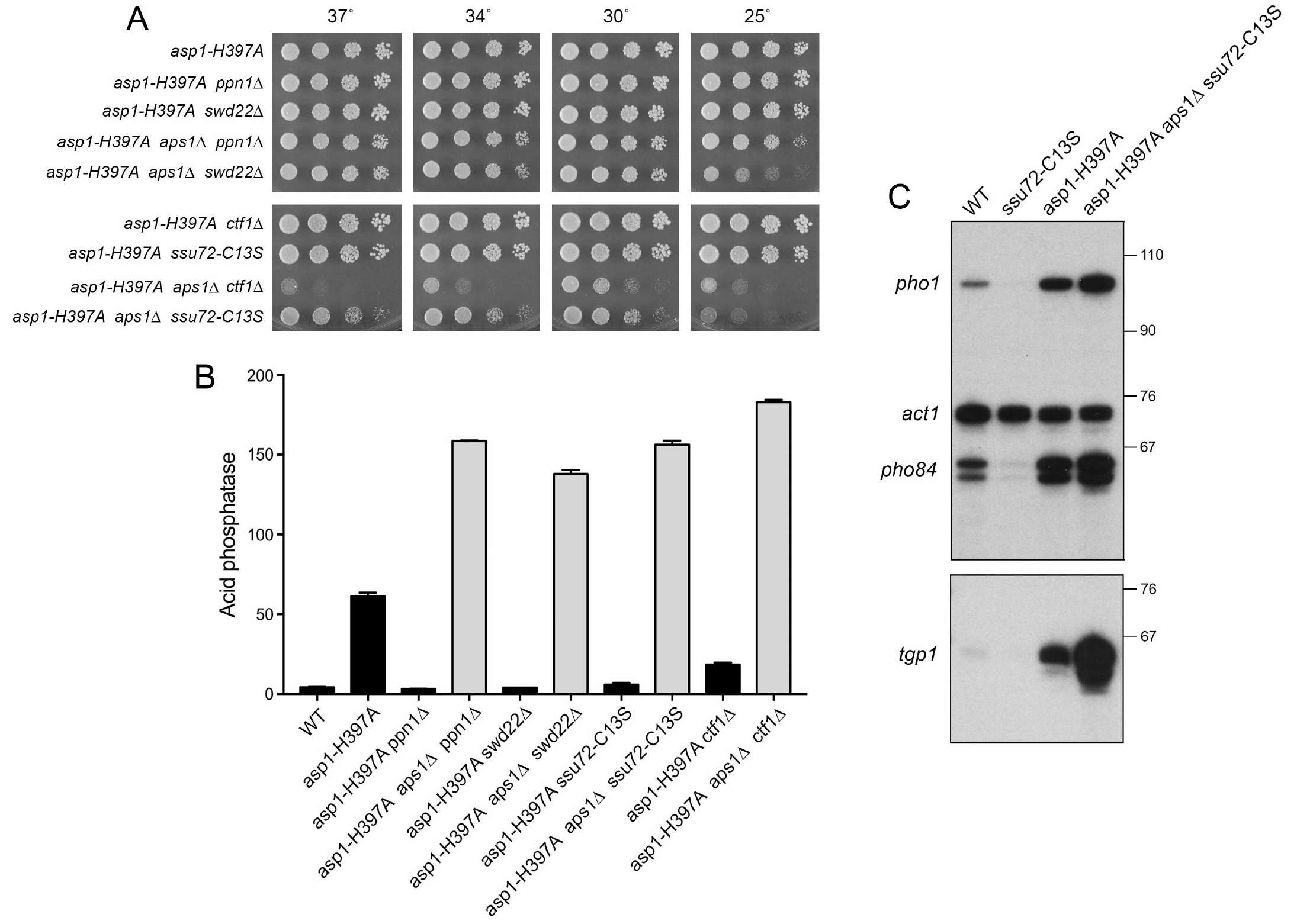
*asp1-H397A aps1*Δ lethality is rescued by *ppn1*Δ, *swd22*Δ, *ssu72-C13S*, and *ctf1*Δ. (A) Strains with the *asp1-H397A* allele in various combinations with *aps1*Δ and CPF subunit mutations were spot tested for growth at the temperatures specified. (B) The indicated strains were grown in liquid culture at 30°C and assayed for acid phosphatase activity. Each datum in the bar graph is the average of assays using cells from at least three independent cultures ± SEM. (C) Total RNA from fission yeast cells with the indicated genotypes was analyzed by reverse transcription primer extension using a mixture of radiolabeled primers complementary to the *pho84*, *act1*, and *pho1* mRNAs (Top panel) or the *tgp1* mRNA (Bottom panel). The reaction products were resolved by denaturing PAGE and visualized by autoradiography. The positions and sizes (nt) of DNA markers are indicated on the right. The result in B and C show that inactivation of the Asp1 and Aps1 pyrophosphatases additively de-represses *pho1*, *pho84*, *and tgp1* expression in a manner that is not prevented by CPF mutations.

### Altered phosphate homeostasis after rescue of *asp1-H397A aps1*Δ by CPF mutants

The rescue of *asp1-H397A aps1*Δ synthetic lethality by *ppn1*Δ, *swd22*Δ, *ssu72-C13S*, and *ctf1*Δ affords an opportunity to gauge the effect of inactivating both IPP pyrophosphatases on *pho1* expression in phosphate-replete cells. The instructive findings were that: (i) Pho1 levels were significantly higher (2.2 to 3-fold) in every one of the four viable *asp1-H397A aps1*Δ strains than in the *asp1-H397A* single-mutant (Fig. 5B); and (ii) adding the *aps1*Δ allele to the *asp1-H397A* double-mutants with *ppn1Δ, swd22*Δ, *ssu72-C13S*, and *ctf1*Δ overrode the antagonistic effects of the CPF mutants on *pho1* de-repression by *asp1-H397A* (Fig. 5B). We conclude that inactivation of the Asp1 and Aps1 pyrophosphatases additively de-represses *pho1* expression in a manner that is not prevented by CPF mutations. The implication is that precocious termination of *prt* lncRNA synthesis is especially sensitive to increased IP8 in *asp1-H397A aps1*Δ cells.

Primer extension analysis of *pho1* mRNA levels in wild-type, *ssu72-C13S*, *asp1-H397A*, and *asp1-H397A aps1Δ ssu72-C13S* triple mutant cells affirmed the acid phosphatase results, i.e., that *pho1* was hyper-repressed by *ssu72-C13S*, de-repressed by *asp1-H397A*, and additively de-repressed in the rescued *asp1-H397A aps1Δ ssu72-C13S* background (Fig. 5C). The same was true of the *pho84* and *tgp1* genes of the phosphate regulon (Fig. 5C).

### Lethality of *dis2*Δ *ssu72-C13S* is rescued by *asp1-H397A*

The synthetic lethality accompanying simultaneous inactivation of any of seven pairwise combinations of CPF subunits and/or Rhn1 (these being *ctf1*Δ *ppn1*Δ, *ctf1*Δ *swd22*Δ, *ppn1*Δ *rhn1*Δ, *swd22*Δ *rhn1*Δ, *ppn1*Δ *ssu72-C13S*, *swd22*Δ *ssu72-C13S*, and *dis2*Δ *ssu72-C13S*; ref. 12) is posited to be the consequence of a severe termination defect impacting the expression of essential *S. pombe* genes. We asked whether such a lethal termination defect in a double-mutant might be ameliorated by a third mutation that exerts an opposite effect, e.g., the *asp1-H397A* allele that we hypothesize promotes precocious termination of the lncRNAs that control phosphate homeostasis. To test this idea, we crossed viable double-mutants of *asp1-H397A* plus *ctf1*Δ, *rhn1*Δ and *ssu72-C13S* (Fig. 2A,C) with differentially marked single mutants *ppn1*Δ, *swd22*Δ, and *dis2*Δ and then screened by random spore analysis for viable triple mutants. Only in the cross of *asp1-H397A ssu72-C13S* with *dis2*Δ did we recover a viable *asp1-H397A dis2*Δ *ssu72-C13S* triple mutant. The *asp1-H397A dis2*Δ *ssu72-C13S* strain grew slower than the *asp1-H397A ssu72-C13S* double mutant on YES agar at 30°C, 34°C and 37°C (as gauged by colony size) and displayed a tight *cs* defect at 25°C and 20°C (Fig. 6A). With respect to Pho1 expression, we saw that the *asp1-H397A dis2*Δ *ssu72-C13S* triple-mutant phenocopied the *asp1-H397A ssu72-C13S* double mutant in erasing the de-repression of Pho1 caused by *asp1-H397A* (Fig. 6B).

**Figure 6.**
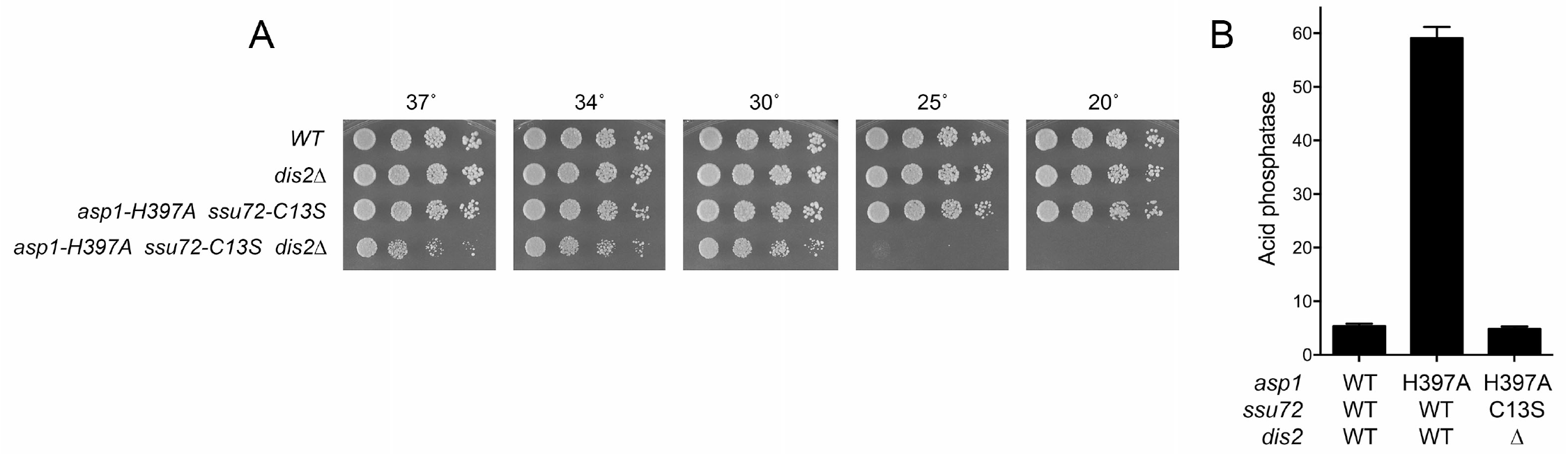
Lethality of *dis2*Δ *ssu72-C13S* is rescued by *asp1-H397A*. (A) Strains with the indicated genotypes were spot tested for growth at the temperatures specified. (B) The indicated strains were grown in liquid culture at 30°C and assayed for acid phosphatase activity.

### Genetic interactions of Asp1 and Aps1 with the Pol2 CTD impact phosphate homeostasis

The de-repressive effect of *asp1-H397A* on *pho1* expression and its genetic reliance on CPF subunits and Rhn1 reported here is similar to the CPF/Rhn1-dependent de-repression of *pho1* observed in *rpb1-CTD-S7A* cells (12). To address whether the de-repression elicited by *S7A* depends on IPP status, we constructed a *rpb1-CTD-S7A asp1-D333A* double-mutant and found that the de-repression of Pho1 by *S7A* was erased when the IPP kinase activity was crippled (Fig. 7A). In the opposite vein, the hyper-repression of *pho1* in *asp1*Δ and *asp1-D333A* cells mimics that seen in *rpb1-CTD-T4A* cells (12, 13). A doubly mutated *rpb1-CTD-(T4A-S7A)* strain maintains a hyper-repressed *pho1* status, implying that the negative effects of *T4A* on *prt* termination win out over the precocious *prt* termination elicited by *S7A* (13). A pertinent question is whether *T4A* exerts a similar effect on the *pho1* de-repression seen in *asp1-H397A* cells.

**Figure 7.**
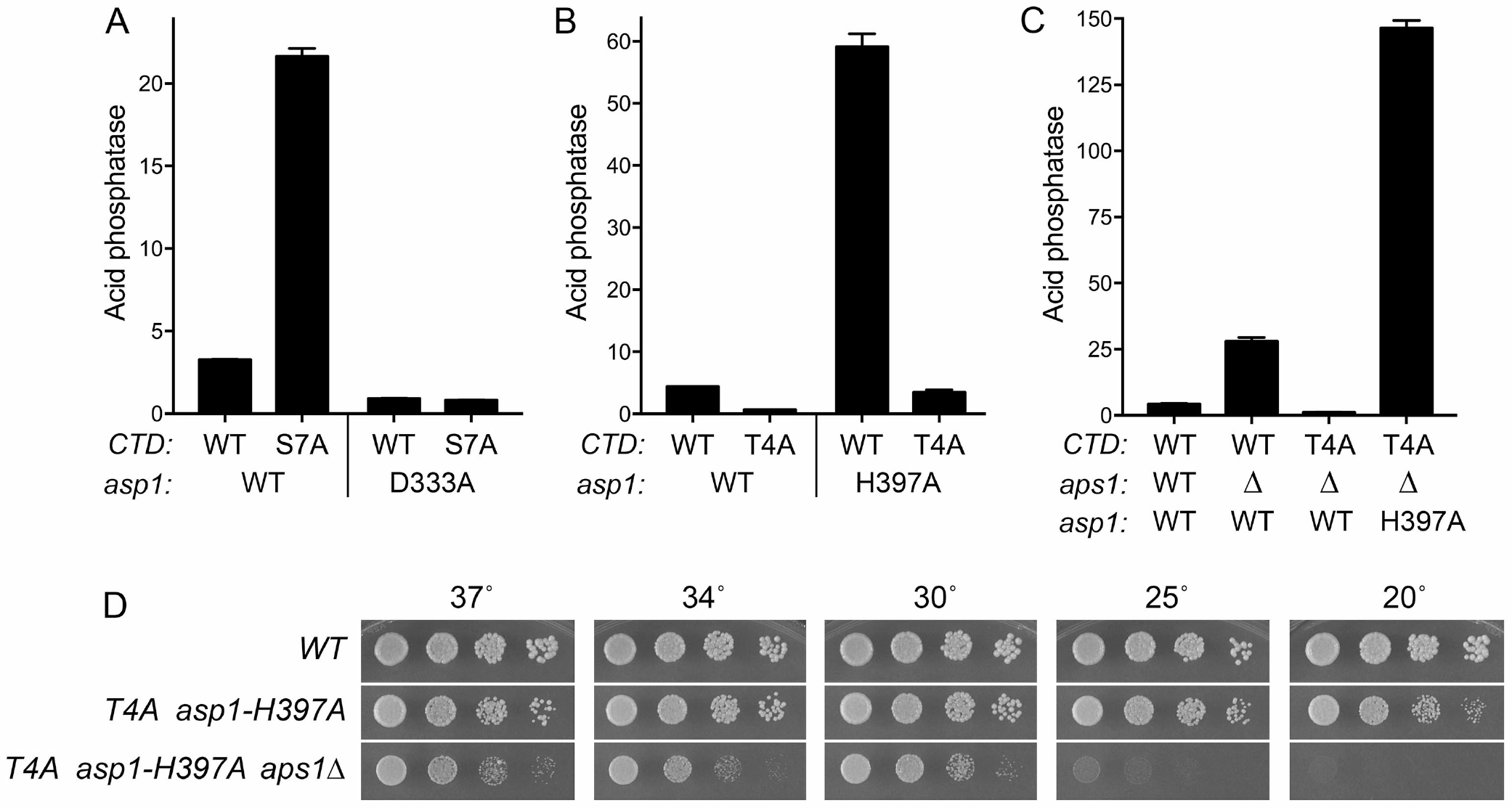
Genetic interactions of Asp1 and Aps1 with the Pol2 CTD impact phosphate homeostasis. (A-C) Strains with the indicated genotypes were grown in liquid culture at 30°C and assayed for acid phosphatase activity. (A) The results show that de-repression of Pho1 by CTD-S7A requires the IPP kinase activity of Asp1. (B) The results show that CTD Thr4 is necessary for de-repression of Pho1 by the *asp1* pyrophosphatase-dead H397A allele. (C) The results show that: (i) CTD Thr4 is necessary for de-repression of Pho1 by *aps1*Δ; and (ii) simultaneous inactivation of Asp1 and Aps1 pyrophosphatases overrides the hyper-repression of Pho1 by *CTD-T4A*. (D) Strains with the indicated genotypes were spot tested for growth on YES agar at the temperatures specified.

We addressed this issue by constructing an *asp1-H397A CTD-T4A* strain; this double mutant thrived at 25°C, 30°C, and 37°C, but was slow growing at 20°C (Fig. 7D). The *CTD-T4A* allele eliminated the large increase in Pho1 activity caused by *asp1-H397A* (Fig. 7B). Similarly, *CTD-T4A* negated the de-repression of Pho1 by *aps1*Δ (Fig. 7C). These results indicate that the precocious *prt* termination prompted by loss of the Asp1 or Aps1 pyrophosphatase activities *per se*, or loss of the Ser7 “letter” of the CTD code, depends stringently on the Thr4 letter (either via the Thr4-PO_4_ mark or the Thr4 hydroxyl group).

It was instructive that the synthetic lethality of the *asp1-H397A aps1*Δ double-mutant was rescued by *CTD-T4A* (Fig. 7D) and that adding *aps1*Δ to the *asp1-H397A CTD-T4A* double-mutant overrode the repressive effect of *T4A* and resulted in 2.5-fold greater Pho1 activity in the *asp1-H397A asp1Δ CTD-T4A* triple-mutant (Fig. 7C) *vis-à-vis* the *asp1-H397A* single mutant (Fig. 7B).

### Synthetic lethalities and their suppression connect IPP status to the Pol2 CTD

Whereas neither *asp1-H397A* nor *CTD-S7A* affects growth of *S. pombe* at 30°C, we were unable to recover an *asp1-H397A CTD-S7A* double-mutant via mating and sporulation, nor were we able to recover an *aps1Δ CTD-S7A* double mutant. These findings signify that IPP accumulation (via inactivation of IPP pyrophosphatases) and loss of the CTD Ser7 mark are synthetically lethal. To query if the synthetic lethality of *asp1-H397A* or *aps1*Δ plus *S7A* mutations reflects a toxic level of precocious termination, we attempted to circumvent the lethality by introduction of mutations in CPF subunits and Rhn1. The striking findings were that: (i) the synthetic lethalities were reversed across the board by *ctf1*Δ, *ssu72-C13S*, *swd22*Δ, *ppn1*Δ, *dis2*Δ, and *rhn1*Δ; and (ii) the rescued triple-mutants grew well at 30°C, 34°C, and 37°C (Fig. 8A,B).

**Figure 8.**
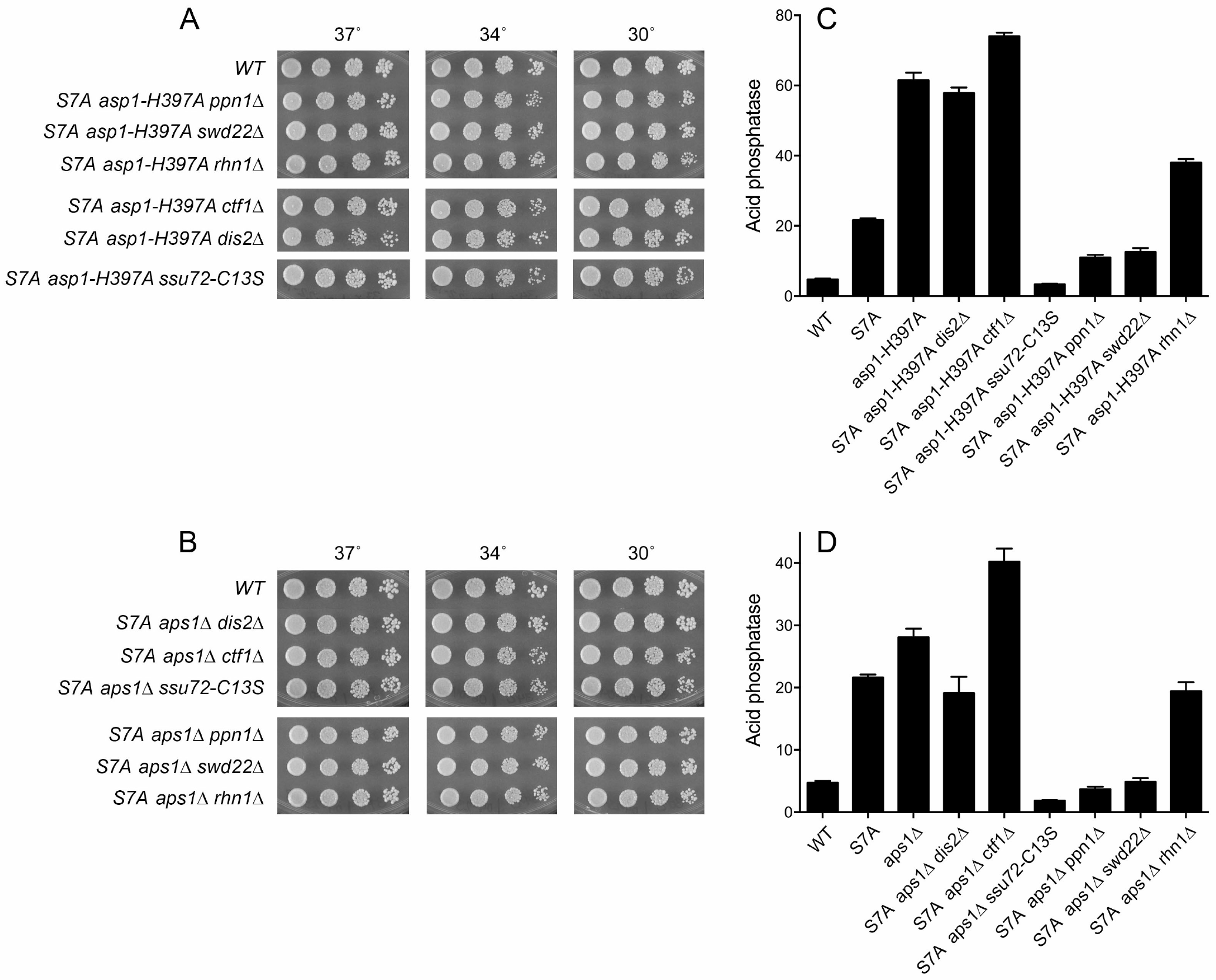
Synthetic lethalities and their suppression connect IPP status to the Pol2 CTD. (A and B) Strains with the indicated genotypes were spot tested for growth at the temperatures specified. The results show that the synthetic lethality of *asp1-H397A* or *aps1*Δ plus *S7A* mutation was reversed by *ctf1*Δ, *ssu72-C13S*, *swd22*Δ, *ppn1*Δ, *dis2*Δ, and *rhn1*Δ. (C and D) The indicated strains were grown in liquid culture at 30°C and assayed for acid phosphatase activity.

We proceeded to gauge Pho1 expression in the rescued CTD-S7A, IPP pyrophosphatase-defective, CPF/Rhn1 triple-mutants, the key question being whether the de-repressive effects of *S7A* and the *asp1-H397A* or *aps1*Δ alleles together “win out” over the hyper-repressive effects of mutations in CPF subunits or Rhn1. The results were remarkable insofar as: (i) CPF/Rhn1 alleles affected Pho1 similarly in the *S7A asp1-H397A* and *S7A aps1*Δ backgrounds (compare Fig. 8C and D); and (ii) there was a clear distinction between the *dis2*Δ, *ctf1*Δ, and *rhn1*Δ alleles that did not reverse Pho1 de-repression in *S7A asp1-H397A* and *S7A aps1*Δ cells and the *ssu72-C13S*, *ppn1*Δ, and *swd22*Δ alleles that eliminated or severely attenuated Pho1 de-repression (Fig. 8C,D). The abilities of *ssu72-C13S*, *ppn1*Δ, and *swd22*Δ to dominate over *S7A asp1-H397A* and *S7A aps1*Δ (Fig. 8C,D) contrast with the “loser” status of these same alleles with respect to *pho1* expression in *asp1-H397A aps1*Δ cells (Fig. 5B), suggesting that the loss of the two IPP pyrophosphatases exerts a stronger effect on precocious termination than does the loss of one IPP pyrophosphatase in combination with CTD-S7A.

### RNA-seq analysis defines an IPP-responsive gene set

The concordant response of the three fission yeast *PHO* regulon genes to IPP pyrophosphatase-inactivating mutations raised the prospect that other genes might be up-regulated when IPP levels are increased in the *asp1-H397A* and *asp1-H397A aps1Δ ssu72-C13S* genetic backgrounds. To explore this idea, we performed RNA-seq on poly(A)^+^ RNA isolated from wild-type, *asp1-H397A, asp1-H397A aps1Δ ssu72-C13S*, and *ssu72-C13S* cells. cDNAs obtained from three biological replicates (using RNA from cells grown to mid-log phase in YES medium at 30°C) were sequenced for each strain. In 11/12 of the datasets, 96-97% of the reads were mapped to unique genomic loci; in one dataset, 90% of the reads mapped to unique loci (Fig. S3). Read densities (RPKM) for individual genes were highly reproducible between biological replicates (Pearson coefficients of 0.98 to 0.99; Fig. S4). As internal controls, we affirmed that: (i) all of the reads for the *asp1* transcript in the *asp1-H397A* strains had the intended His-to-Ala codon mutation; (ii) all the reads for the *ssu72* transcript in the *ssu72-C13S* strains had the Cys-to-Ser codon mutation; and (iii) there were no reads for the deleted *aps1* coding sequence in the *asp1-H397A aps1Δ ssu72-C13S* strain. A cutoff of ±2-fold change in normalized transcript read level and a corrected p-value of ≤0.05 were the criteria applied to derive initial lists of differentially expressed annotated loci in the mutants versus wild-type. We then focused on differentially expressed coding genes with average normalized read counts across all samples of >100 (DESeq2 baseMean parameter), in order to eliminate the many, mostly non-coding, transcripts that were expressed at very low levels in vegetative cells. The list of genes that were dysregulated by these criteria in one or more of the mutant strains is compiled in Supplemental Table S1.

Figure 9A shows the list of 30 annotated protein-coding genes that were up-regulated in *asp1-H397A* and *asp1-H397A aps1Δ ssu72-C13S* cells, but not in *ssu72-C13S* cells. This gene set comprises a putative IP8-responsive regulon. The “top hits” with respect to fold upregulation included the three known phosphate-regulated genes: *tgp1* (up 21-fold in *asp1-H397A* and 42-fold *asp1-H397A aps1Δ ssu72-C13S*); *pho1* (up 7-fold in *asp1-H397A* and 14-fold *asp1-H397A aps1Δ ssu72-C13S*); and *pho84* (up 4-fold in *asp1-H397A* and 6-fold *asp1-H397A aps1Δ ssu72-C13S*). Thus, the RNA-seq data independently affirms the conclusions from assays of Pho1 enzyme activity, and of *pho1*, *pho84*, and *tgp1* mRNA levels assayed by primer extension, that the *PHO* regulon is coordinately de-repressed when Asp1 pyrophosphatase is defective. Note: it is not the case that the *PHO* genes were overexpressed as a secondary effect of altered expression of other genes involved in *PHO* transcription/homeostasis or of genes encoding the 3’ processing/termination machinery. RNA-seq of *asp1-H397A* and *asp1-H397A aps1Δ ssu72-C13S* cells revealed insignificant effects on the transcripts encoding Pho7 (the transcription factor that drives *pho1*, *pho84*, and *tgp1* mRNA synthesis), the Rpb1 subunit of Pol2, protein kinases Csk1 and Cdk9 (mutations of which de-repress *pho1*), cleavage/polyadenylation factors Ctf1, Cft1, Cft2, Dis2, Iss1, Pfs2, Pla1, Pta1, Ppn1, Rna14, Ssu72, Swd22, Ysh1, Yth1, Pcf11, and termination factors Rhn1, Seb1, and Dhp1.

**Figure 9.**
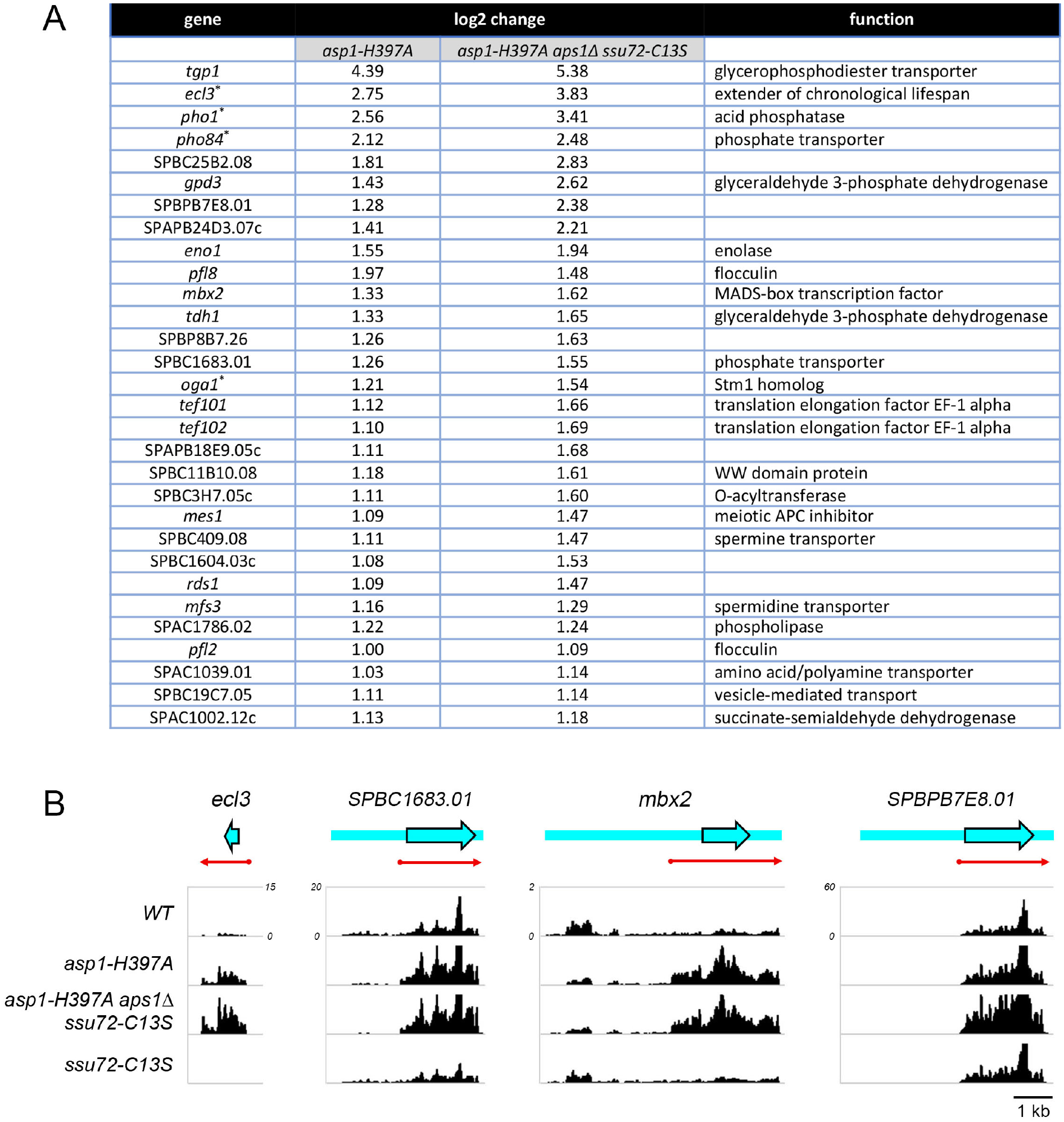
RNA-seq defines an IPP-responsive gene set. (A) List of annotated protein coding genes that were up-regulated at least two-fold in *asp1-H397A* and *asp1-H397A aps1Δ ssu72-C13S* cells. Any genes that were also upregulated two-fold in *ssu72-C13S* cells were excluded. The four genes denoted by asterisks (*pho1*, *pho84*, *ecl3*, and *oga1*) were down-regulated by at least two-fold in the *ssu72-C13S* strain. (B) Strand-specific RNA-seq read densities (counts/base/million, averaged in a 25 nucleotide window) of the indicated *S. pombe* strains are plotted on the *y*-axis as a function of position across the gene loci (*x*-axis). The read densities were determined from cumulative counts of all three RNA-seq replicates for each *S. pombe* strain. The *y*-axis scale for all tracks of each individual gene is shown next to the *WT* track. The common *x*-axis scale is shown on the bottom right. ORFs are indicated by blue arrowheads with black outline and UTRs by blue bars extending from the ORF, shown as they are annotated in Pombase. Red arrows indicate the predicted mRNA as judged from the RNA-seq data.

Another top upregulated hit in the IPP-responsive gene set (Fig. 9A) was *ecl3* (up 7-fold in *asp1-H397A* and 14-fold in *asp1-H397A aps1Δ ssu72-C13S*), which encodes an 89-amino acid protein that extends fission yeast chronological lifespan when overexpressed (31). It is noteworthy that: (i) the *ecl3* gene is located on chromosome II, adjacent to and in opposite orientation to the *prt2* lncRNA gene of the phosphate-regulated *prt2*–*pho84*–*prt*–*pho1* gene cluster (10); and (ii) expression of the *ecl3* paralogs *ecl1* and *ecl2*, which are located at distant sites in the genome (31), is not affected in *asp1-H397A* and *asp1-H397A aps1Δ ssu72-C13S* cells. We speculate that *ecl3* up-regulation by inactivation of IPP pyrophosphatase might reflect its proximity to *prt2*, e.g., that the *prt2* promoter might be bidirectional (*à la* the *nc-tgp1* promoter; 6), driving transcription of the *prt2* lncRNA that regulates *pho84* and an oppositely transcribed lncRNA that interferes with *ecl3* transcription. Consistent with the idea that de-repression of *ecl3* by IPP pyrophosphatase inactivation might be connected to lncRNA transcription termination, the RNA-seq analysis showed that the *ecl3* transcript was downregulated in *ssu72-C13S* cells (by 3-fold), concordant with the downregulation seen in *ssu72-C13S* cells for *pho1* (by 5-fold) and *pho84* (by 5-fold). Annotation of the *ecl3* gene in PomBase (www.pombase.org) has no information other than the dimensions of the open reading frame (ORF, denoted by the blue arrow in Fig. 9B, left panel). Plots of the strand-specific RNA-seq read density across this region in the *WT*, *asp1-H397A*, *asp-H397A aps1*Δ *ssu72-C13S*, and *ssu72-C13S* strains affirmed that *ecl3* expression is upregulated by IPP pyrophosphatase inactivation and downregulated by Ssu72 inactivation (Fig. 9B). Moreover, the read density in the upregulated strains clearly delineates the approximate margins of the *ecl3* mRNA upstream and downstream of the ORF (red arrow in Fig. 9B).

The IPP-responsive gene set also includes SPBC1683.01, which encodes a 573-amino acid putative inorganic phosphate transporter that is 84% identical in primary structure to the 572-amino acid Pho84 protein. The SPBC1683.01 mRNA is annotated in PomBase as a 3923-nucleotide transcript that includes a 1950-nucleotide 5’-UTR (Fig. 9B). Because this long 5’-UTR contains 36 AUG codons upstream of the *bona fide* translation start site of the transporter ORF, we are skeptical of the transcript annotation. Rather, as we showed for *prt2–pho84* (10), we suspect that the ∼2-kb region upstream of the transporter ORF specifies an independently transcribed lncRNA that regulates in *cis* the expression of the downstream SPBC1683.01 phosphate transporter. This idea is supported by the strand-specific RNA-seq read density across this gene in the *WT*, *asp1-H397A*, *asp-H397A aps1*Δ *ssu72-C13S*, and *ssu72-C13S* strains (Fig. 9B), insofar as there is a selective increase in RNA reads over the ORF upon IPP pyrophosphatase inactivation that is accompanied by a decrease in reads over the so-called 5’-UTR. By contrast, Ssu72 inactivation reduced reads over the ORF and tended to even out the read density across the locus (Fig. 9B). The read density in the upregulated strains demarcates the approximate margins of the SPBC1683.01 mRNA (red arrow in Fig. 9B). The results suggest that the putative lncRNA is a read-through transcript.

The *mbx2* gene that is upregulated in *asp1-H397A* and *asp1-H397A aps1Δ ssu72-C13S* cells encodes a 372-amino acid MADS-box transcription factor that positively regulates invasive growth and flocculation by driving the expression of cell surface flocculin proteins (32, 33). The IPP-responsive upregulated gene set includes the Mbx2-regulated flocculin genes *pfl8* and *pfl2* (Fig. 9A). Moreover, the flocculin genes *gsf2* and *pfl3* are upregulated in *asp1-H397A* and *asp1-H397A aps1Δ ssu72-C13S* cells (though they are also upregulated in the *ssu72-C13S* strain). The increased expression of the Mbx2 transcription factor and several flocculin proteins neatly accounts for the findings that *asp1-H397A* cells display a strong flocculation phenotype and hyper-invasive growth *vis-à-vis asp1*^+^ cells (17). It is noteworthy that *mbx2* mRNA is annotated in PomBase as a 6079-nucleotide transcript with a 4059-nucleotide 5’-UTR (Fig. 9B) that contains 69 AUG codons upstream of the translation start site of the Mbx2 ORF. Thus, we speculate that the ∼4-kb region upstream of the Mbx2 ORF specifies a lncRNA that regulates the expression of the Mbx2 mRNA. The RNA-seq read density in the upregulated IPP pyrophosphatase-inactive strains indicates the dimensions of the true *mbx2* mRNA (red arrow in Fig. 9B).

Yet another example of such potential regulation in the IPP-responsive gene set is SPBPB7E8.01, which encodes an uncharacterized 569-amino acid protein. The SPBPB7E8.01 mRNA is annotated as a 4959-nucleotide transcript with a 2698-nucleotide 5’-UTR (Fig. 9B) that contains 41 AUG codons upstream of the translation start site of the SPBPB7E8.01 ORF. Again, we envision that the ∼2.7-kb region upstream of the SPBPB7E8.01 ORF is transcribed independently as a lncRNA that regulates the expression of the SPBPB7E8.01 mRNA. The RNA-seq read densities clearly delineate the margins of the SPBPB7E8.01 mRNA (red arrow in Fig. 9B) and show that it does not include the annotated long 5’-UTR.

In sum, the above genome-wide RNA-seq analysis affirms the conclusions from the preceding gene-specific experiments (at RNA and protein level) that the fission yeast *PHO* regulon is de-repressed by IPP pyrophosphatase inactivation and it delineates a novel IPP-responsive gene regulon, many members of which are known to be (e.g. *tgp1*, *pho1*, *pho84*), or are likely to be, under repressive control by an upstream flanking lncRNA.

### Transcriptome analysis of *aps1*Δ accords with that of *asp1-H397A*

To gauge the effect of loss of the Aps1 pyrophosphatase activity *per se* on gene expression, we performed RNA-seq of poly(A)^+^ RNA isolated from wild-type and *aps1*Δ cells, sequencing cDNAs obtained from three biological replicates (see Figs. S5 and S6 for compilation of read counts and reproducibility of read densities between biological replicates). Figure 10 shows the list of 19 annotated protein-coding genes that were up-regulated in *aps1*Δ cells. Nine of these 19 genes (denoted by asterisks in Fig. 10) were upregulated in *asp1-H397A* cells (Fig. 9A). The co-regulated gene set includes the three members of the *PHO* regulon and the *prt2*-adjacent *ecl3* gene. The *tgp1*, *ecl3*, *pho1*, and *pho84* transcripts were increased by 7-fold, 4-fold, 3-fold, and 3-fold in *aps1*Δ cells. We also observed upregulation of *mbx2* and of several of the flocculin genes under its transcriptional control. The concordant overlapping set of upregulated genes in two different genetic backgrounds in which distinct IPP pyrophosphatase activities are missing fortifies the case for an IPP-responsive transcriptional program in fission yeast.

**Figure 10.**
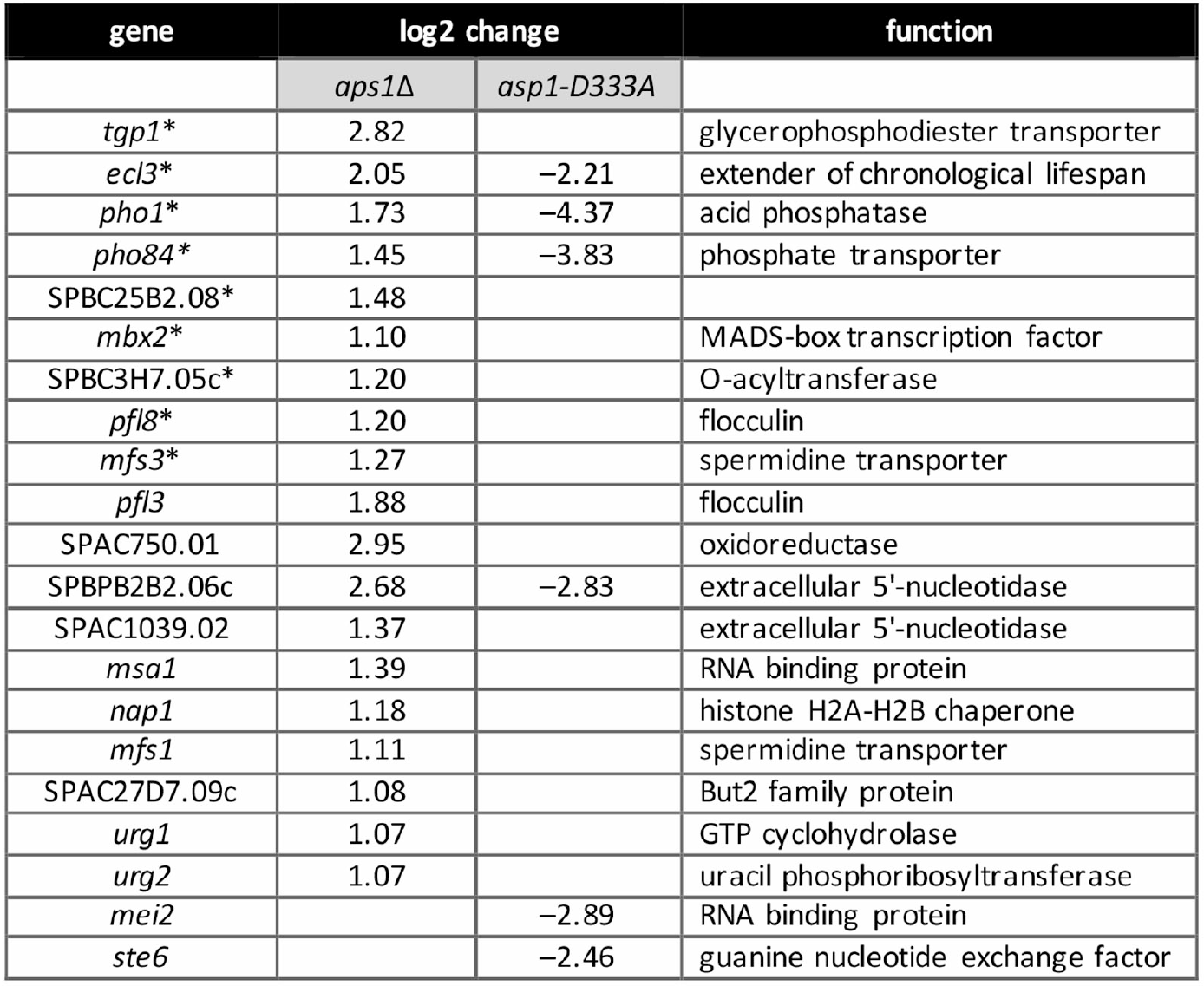
Transcriptome profile of *aps1*Δ and *asp1-D333A* cells. List of 19 annotated protein-coding genes that were up-regulated at least two-fold in *aps1*Δ cells and 6 annotated protein-coding genes that were downregulated at least four-fold in *asp1-D333A* cells. Nine genes that were coordinately upregulated in *aps1*Δ and *asp1-H397A* IPP pyrophosphatase-deficient cells are denoted by asterisks.

### Transcriptome analysis of *asp1*-*D333A*

If, as we hypothesize, increased levels of IP8 in the IPP pyrophosphatase-defective strains is responsible for the observed increases in a limited set of mRNAs, then we might expect to see opposite effects on some of these IPP-responsive transcripts when IP8 synthesis is precluded by the kinase-defective *asp1-D333A* allele. RNA-seq analysis of poly(A)^+^ RNA from *asp1-D333A* cells identified six protein-coding genes that were down-regulated by at least 4-fold compared to the wild-type control (Fig. 10), four of which correspond to transcripts that were upregulated in *aps1*Δ and/or *asp1-H397A* cells. [As an internal control, we affirmed that all of the reads for the *asp1* transcript in the *asp1-D333A* strains had the intended Asp-to-Ala codon mutation.] The two top hits in the down-regulated *asp1-D333A* gene set were *pho1* and *pho84*, with decrements of 20-fold and 14-fold, respectively. The *ecl3* transcript was reduced 5-fold in *asp1-D333A* cells. SPBPB2B2.06c encoding a putative extracellular 5’ nucleotidase was 7-fold down in *asp1-D333A* cells. Of the other *aps1*Δ or *asp1-H397A* up-regulated genes, only *mfs3* was appreciably down-regulated in *asp1-D333A* cells (by 3-fold). Thus, there appear to be two classes of IPP-responsive genes: those that require IP8 for normal expression and are overexpressed when IP8 levels are elevated above normal (e.g., the *PHO* genes); and those that do not require IP8 for normal expression but are overexpressed when IP8 levels are elevated.

## DISCUSSION

Fission yeast phosphate acquisition genes *pho1*, *pho84*, and *tgp1* are repressed during growth in phosphate-rich medium by the transcription in *cis* of upstream flanking lncRNAs. This transcription interference phenomenon is tuned by the phosphorylation status of the RNA polymerase II CTD and the RNA 3’ processing/termination factors CPF and Rhn1 in a manner whereby genetic changes that elicit precocious termination of lncRNA synthesis lead to de-repression of downstream mRNA expression (e.g., *rpb1-CTD-S7A*) and changes that diminish lncRNA termination hyper-repress mRNA expression (e.g., *rpb1-CTD-T4A*, CPF subunit mutations, *rhn1*Δ). In the present study, we show that fission yeast phosphate homeostasis is subject to metabolite control by inositol pyrophosphates (IPPs), exerted via the 3’ processing/termination machinery and the Pol2 CTD.

The genetic evidence implicating IPPs, particularly IP8, as a new player in the interactome of the CTD code with 3’ processing and termination, stems from the coherent set of biochemical phenotypes and mutational synergies elicited by manipulations of the kinase and pyrophosphatase enzymes that convert IP7 to IP8 and *vice versa*. It is already established that: (i) inactivation of the Asp1 kinase component eliminates IP8 and increases intracellular IP7; and (ii) inactivation of the Asp1 pyrophosphatase module increases IP8 without much effect on IP7 (19). Here we find that increasing IP8 via IPP pyrophosphatase mutation de-represses the *PHO* regulon. Focusing on *pho1* de-repression, we show that increased IP8 (or increased IP8:IP7 ratio) leads to precocious termination during *prt* lncRNA synthesis, and that *pho1* de-repression depends on CPF subunits, Rhn1, and the Thr4 letter of the CTD code. Conversely, a failure to synthesize IP8 (via IPP kinase mutation) results in *pho1* hyper-repression, thereby phenocopying mutations of CPF subunits, Rhn1, and CTD-T4A. These findings suggest that IP8 enhances the responsiveness of elongating Pol2 to the action of cleavage/polyadenylation and termination factors. Indeed, the de-repression of *pho1* caused by the mutating CTD Ser7 to alanine depends on the Asp1 kinase that generates IP8. Moreover, the synthetic lethality of *asp1*Δ (no IP8) with mutations of CPF subunits Ppn1, Swd22, and Ssu72 argues that IP8 plays an important role in promoting essential 3’ processing/transcription termination events in fission yeast, albeit in a manner that is genetically redundant to CPF.

Too much IP8 is apparently toxic to fission yeast, insofar as simultaneous inactivation of the *asp1* and *aps1* IPP pyrophosphatase enzymes is lethal. That this lethality is tied to RNA 3’ processing/termination and the CTD code is established by the finding that *asp1-H397A aps1*Δ inviability is suppressed by mutations of CPF subunits Ppn1, Swd22, Ssu72, and Ctf1 and by CTD-T4A. In the same vein, increasing IP8 by inactivating the Asp1 pyrophosphatase is lethal in combination with CTD-S7A, implying that these changes additively promote precocious termination to the point that it becomes toxic. Here, too, the lethality of *asp1-H397A CTD-S7A* is suppressed by mutations of CPF subunits and termination factor Rhn1. Lastly, the yin-yang relationship of IPP pyrophosphatase mutations (that elicit precocious termination) and CPF mutations (that diminish termination) is cemented by the finding that the synthetic lethality of pairwise mutations of CPF subunits Dis2 and Ssu72 is suppressed by *asp1-H397A* allele that increases IP8.

Whereas IPPs have been implicated in a broad range of physiological processes (34, 35), the present study is, to our knowledge, unique in forging a link between perturbations of IPP dynamics and RNA 3’ processing/transcription termination. The immediate target(s) of IP8 in eliciting the effects we describe here are as yet unknown. Two distinct themes have been invoked previously to account for IPP effects, whereby IPPs either: (i) bind to target proteins and allosterically enhance or inhibit their functions; or (ii) serve as phosphate donors for transfer of an IPP β-phosphate to a previously mono-phosphorylated protein target leading to a covalent pyrophosphate modification of the target (34–37). Potential mechanisms for IP8 to promote 3’-processing/termination include: (i) allosteric binding or covalent modification of the processing/termination machinery so as to enhance its action on elongating Pol2; (ii) allosteric interaction with, or covalent modification of Pol2 to make it more responsive to the processing/termination machinery. Given the connections established here between IP8 and the Pol2 CTD phospho-site code, an enticing speculation is that increased IP8 levels elicit pyrophosphorylation of the phospho-CTD. There is precedent for allosteric interactions of IPPs with protein kinases that result in inhibition of kinase activity (36). The fission yeast Csk1 kinase is critical for repression of *pho1* under phosphate-replete conditions, i.e., *pho1* is de-repressed in a *csk1*Δ strain (15) in a manner that depends on Ssu72 and Rhn1 (12). Because the *csk1*Δ and IPP pyrophosphatase-dead mutations have overlapping phosphate homeostasis phenotypes, we reasoned that if the effect of increasing IP8 was transduced via inhibition of Csk1 kinase, then there would be no mutational synergy in a *csk1*Δ *asp1-H397A* double mutant. However, this model was vitiated by our finding that *csk1*Δ and *asp1-H397A* were synthetically lethal (not shown).

The SPX protein domain has been identified as an inositol polyphosphate/inositol pyrophosphate sensor found in plant and budding yeast proteins involved in phosphate homeostasis (38, 39). Fission yeast have six annotated SPX domain-containing proteins (www.pombase.org). However, an SPX domain is not associated with any fission yeast proteins implicated in 3’ processing or transcription termination.

The ENTH protein domain involved in membrane traffic binds to inositol polyphosphates (40). The α-helical fold of the ENTH domain is very similar to that of the VHS domain (41) and to that of the Pol2 CTD-interaction domain (CID) found in several components of the mRNA transcription/processing apparatus (42). Fission yeast CID-containing proteins include CPF subunit Pcf11 and termination factors Rhn1 and Seb1. CID proteins have not (to our knowledge) been reported to bind IPPs. Our inspection of the structure of *S. pombe* Seb1 (43) suggests that it may have a counterpart of the lysine-rich surface motif responsible for inositol polyphosphate binding in the ENTH domain (40). Seb1 interacts physically with the fission yeast CPF complex (43). It is conceivable that IPP interaction with Seb1 (or one of the other fission yeast CID proteins) enhances 3’-processing/termination.

Finally, transcriptional profiling of IPP pyrophosphatase-defective *S. pombe* strains delineated what we construe to be an IP8-responsive regulon composed of genes that are overexpressed when IP8 levels are increased. The IP8-responsive gene set includes the phosphate acquisition regulon, which is repressed by upstream lncRNA synthesis, as well as other genes that are plausible candidates for similar lncRNA control.

## Acknowledgements

We thank Dr. Ursula Fleig (Heinrich Heine Universität) for providing the fission yeast *asp1-D333A* and *asp1-H397A* strains and the expression plasmids for the Asp1 kinase domain and pyrophosphatase domain. This work was supported by NIH grants R01-GM52470 and R35-GM126945.

## Data Deposition

The RNA-seq data in this publication have been deposited in NCBI’s Gene Expression Omnibus and are accessible through GEO Series accession numbers GSE12755 and (https://www.ncbi.nlm.nih.gov/geo/query/acc.cgi?acc=GSE127550) and GSE131237 (https://www.ncbi.nlm.nih.gov/geo/query/acc.cgi?acc=GSE131237).

