## Supplemental Material for "Inositol pyrophosphates impact phosphate homeostasis via modulation of RNA 3’ processing and transcription termination"

Supplemental Figures S1, S2, S3, S4, S5, S6, S7,

Supplemental Tables S1 and S2 (provided as separate Excel files)

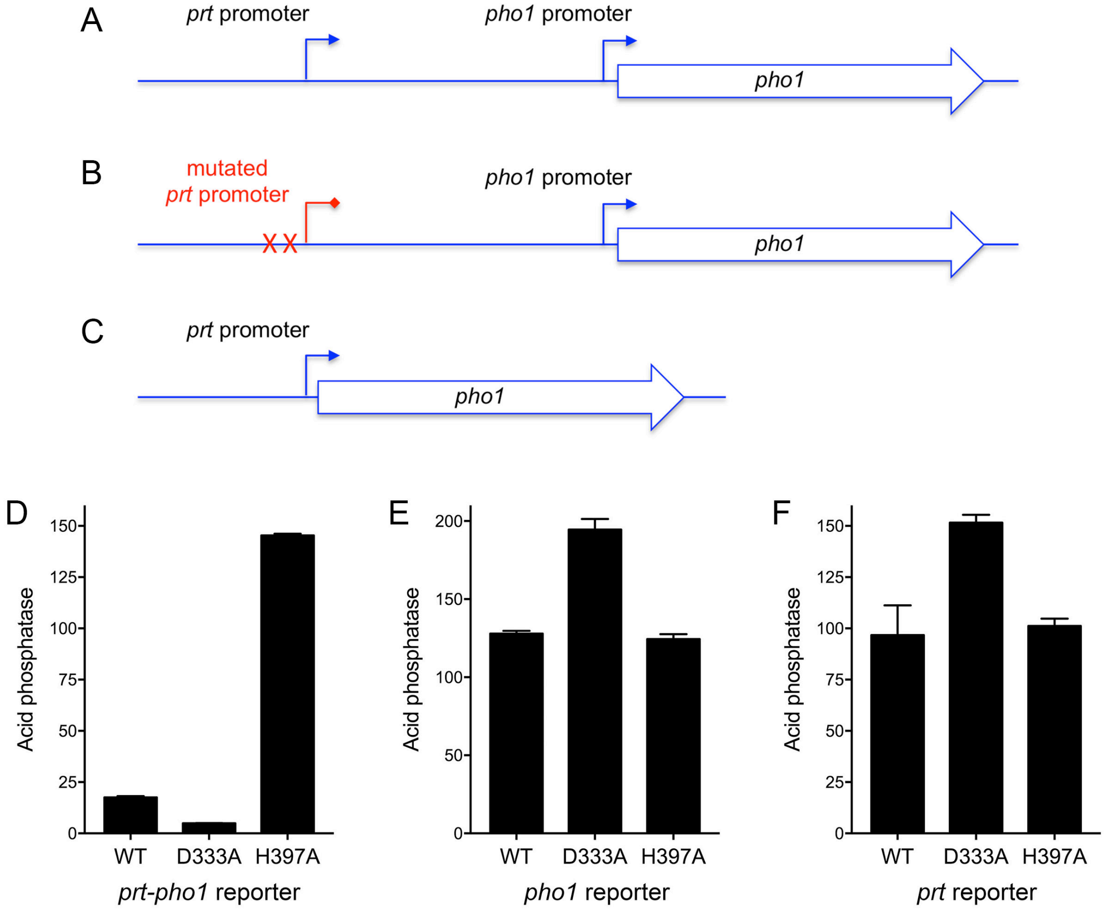


Figure S1. **Do Asp1 perturbations affect the *prt* lncRNA or *pho1* mRNA promoters?** (A) Schematic of the plasmid-borne *prt-pho1* reporter in which pho1 expression is repressed by *prt* lncRNA transcription. (B) A reporter of *pho1* promoter activity in which *prt* lncRNA transcription is abolished by mutations (indicated by X) in the HomolD and TATA box elements in the *prt* promoter as described (8). (C) A reporter of *prt* promoter activity in which the *prt* promoter directly drives transcription of the *pho1* gene. (D,E,F) The indicated reporter plasmids were transfected into *asp1*$^{+}$ (WT), *asp1-D333A* (kinase-dead), or *asp1-H397A* (pyrophosphatase-dead) strains in which the chromosomal *pho1* locus was deleted (10). Transformants were selected and single colonies of individual transformants were pooled (>20) and grown in plasmid-selective liquid medium to $A_{600}$ of 0.5-0.8. Aliquots were harvested by centrifugation for acid phosphatase activity measurements. Each datum in the bar graph is the average of assays using cells from three independent cultures ± SEM.

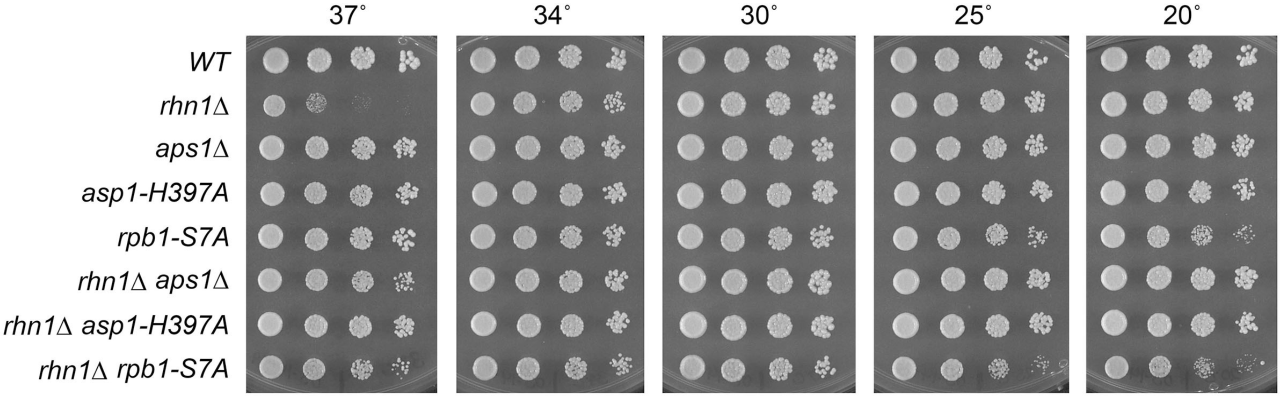


Figure S2. **The *ts* growth defect of *rhn1*Δ is suppressed by inactivation of IPP pyrophosphatases and CTD-S7A**. *S. pombe* strains with the genotypes indicated at left were inoculated in YES broth and grown at 30°C. Exponentially growing cultures were adjusted to $A_{600}$ of 0.1 and aliquots (3 μl) of serial 5-fold dilutions were spotted on YES agar and then incubated at the temperatures specified.

| Sample | Total Paired Reads | Mapped Reads |
|---|---|---|
| *WT* (1) | 32,971,120 | 31,701,973 (96%) |
| *WT* (2) | 29,319,196 | 28,367,274 (97%) |
| *WT* (3) | 27,207,850 | 26,332,834 (97%) |
| *ssu72-C13S* (1) | 30,789,188 | 29,648,320 (96%) |
| *ssu72-C13S* (2) | 28,737,729 | 27,757,868 (97%) |
| *ssu72-C13S* (3) | 30,792,948 | 27,669,979 (90%) |
| *asp1-H397A* (1) | 27,947,318 | 27,079,708 (97%) |
| *asp1-H397A* (2) | 30,981,782 | 29,833,174 (96%) |
| *asp1-H397A* (3) | 28,254,348 | 27,415,167 (97%) |
| *asp1-H397A asp1Δ ssu72-C13S* (1) | 28,342,783 | 27,337,584 (96%) |
| *asp1-H397A asp1Δ ssu72-C13S* (2) | 31,180,683 | 30,108,674 (97%) |
| *asp1-H397A asp1Δ ssu72-C13S* (3) | 26,054,969 | 25,167,313 (97%) |

Figure S3. **RNA-seq read counts for triplicate biological replicates**

| Sample pairs | Pearson Coefficient |
|---|---|
| *WT* (1) vs (2) | 0.989 |
| *WT* (2) vs (3) | 0.990 |
| *WT* (1) vs (3) | 0.989 |
| *ssu72-C13S* (1) vs (2) | 0.989 |
| *ssu72-C13S* (2) vs (3) | 0.988 |
| *ssu72-C13S* (1) vs (3) | 0.988 |
| *asp1-H397A* (1) vs (2) | 0.988 |
| *asp1-H397A* (2) vs (3) | 0.980 |
| *asp1-H397A* (1) vs (3) | 0.981 |
| *asp1-H397A asp1Δ ssu72-C13S* (1) vs (2) | 0.983 |
| *asp1-H397A asp1Δ ssu72-C13S* (2) vs (3) | 0.987 |
| *asp1-H397A asp1Δ ssu72-C13S* (1) vs (3) | 0.982 |

Figure S4. **RNA-seq data reproducibility between biological replicates**

| Sample | Total Paired Reads | Mapped Reads |
|---|---|---|
| *WT* (4) | 21449706 | 20980489 (98%) |
| *WT* (5) | 23888710 | 23365656 (98%) |
| *WT* (6) | 22670827 | 22083962 (97%) |
| *aps1Δ* (1) | 22637872 | 21614517 (95%) |
| *aps1Δ* (2) | 21756873 | 21114730 (97%) |
| *aps1Δ* (3) | 22454601 | 21932787 (98%) |
| *asp1-D333A* (1) | 22976308 | 22449419 (98%) |
| *asp1-D333A* (2) | 25347559 | 24797369 (98%) |
| *asp1-D333A* (3) | 25475939 | 24908694 (98%) |

Figure S5. **RNA-seq read counts for triplicate biological replicates**

| Sample pairs | Pearson Coefficient |
|---|---|
| *WT* (4) vs (5) | 0.976 |
| *WT* (5) vs (6) | 0.987 |
| *WT* (4) vs (6) | 0.978 |
| *aps1Δ* (1) vs (2) | 0.983 |
| *aps1Δ* (2) vs (3) | 0.988 |
| *aps1Δ* (1) vs (3) | 0.983 |
| *asp1-D333A* (1) vs (2) | 0.981 |
| *asp1-D333A* (2) vs (3) | 0.979 |
| *asp1-D333A* (1) vs (3) | 0.977 |

Figure S6. **RNA-seq data reproducibility between biological replicates**

| **Strain** | **Relevant genotype** | **Source (reference)** |
|---|---|---|
| JS77<br>JS78 | *ade6-m216 leu1-32 ura4-D18 his3-D1* $h^-$<br>*ade6-m210 leu1-32 ura4-D18 his3-D1* $h^+$ | Pei et al. 2006 Mol Cell Biol 26: 777-788. |
| AS 2058 | *rpb1-T4A$_{29}$::natMX* | Sanchez et al. 2018 (12) |
| AS 1830 | *rpb1-S7A$_{29}$::natMX* | Sanchez et al. 2018 (12) |
| AS 1789 | *ctf1Δ::kanMX* | Sanchez et al. 2018 (12) |
| AS 1823 | *ctf1Δ::ura4$^+$* | Sanchez et al. 2018 (12) |
| AS 386 | *ssu72-C13S::kanMX* | Schwer et al. 2015 (7) |
| AS 612 | *ssu72-C13S::natMX* | Sanchez et al. 2018 (12) |
| AS 1838 | *rhn1Δ::kanMX* | Sanchez et al. 2018 (12) |
| AS 1911 | *rhn1Δ::natMX* | Sanchez et al. 2018 (12) |
| AS 2003 | *rhn1Δ::hygMX* | Sanchez et al. 2018 (12) |
| AS 1940 | *dis2Δ::ura4$^+$* | R. Fisher |
| AS 1979 | *ppn1Δ::hygMX* | Sanchez et al. 2018 (12) |
| AS 1980 | *swd22Δ::hygMX* | Sanchez et al. 2018 (12) |
| AS 1929 | *[prt2-pho84-prt-pho1]Δ::hygMX* | Garg et al. 2018 (10) |
| AS2225 | *[prt2-pho84-prt-pho1]Δ::hygMX asp1-D333A::natMX* | This study |
| AS2226 | *[prt2-pho84-prt-pho1]Δ::hygMX asp1-H397A::natMX* | This study |
| AS1886 | *asp1Δ::kanMX* | This study |
| AS2183 | *asp1Δ::natMX* | This study |
| AS1884 | *aps1Δ::kanMX* | This study |
| AS1917 | *aps1Δ::natMX* | This study |
| AS2287 | *aps1Δ::hygMX* | This study |
| AS2180 | *asp1-D333A::kanMX* | U. Fleig |
| AS2184 | *asp1-D333A::natMX* | This study |
| AS2181 | *asp1-H397A::kanMX* | U. Fleig |
| AS2185 | *asp1-H397A::natMX* | This study |
| AS2211 | *aps1Δ::natMX asp1Δ::kanMX* | This study |
| AS2212 | *aps1Δ::natMX asp1-D333A::kanMX* | This study |
| AS2152 | *aps1Δ::natMX ctf1Δ::kanMX* | This study |
| AS2153 | *aps1Δ::natMX rhn1Δ::kanMX* | This study |
| AS2154 | *aps1Δ::natMX ssu72-C13S::kanMX* | This study |
| AS2155 | *aps1Δ::natMX dis2Δ::ura4$^+$* | This study |
| AS2157 | *aps1Δ::natMX ppn1Δ::hygMX* | This study |
| AS2158 | *aps1Δ::natMX swd22Δ::hygMX* | This study |
| AS2186 | *asp1-H397A::kanMX ctf1Δ::ura4$^+$* | This study |
| AS2187 | *asp1-H397A::kanMX dis2Δ::ura4$^+$* | This study |
| AS2188 | *asp1-H397A::kanMX rhn1Δ::natMX* | This study |
| AS2189 | *asp1-H397A::kanMX ssu72-C13S::natMX* | This study |
| AS2190 | *asp1-H397A::kanMX ppn1Δ::hygMX* | This study |
| AS2191 | *asp1-H397A::kanMX swd22::hygMX* | This study |
| AS2196 | *asp1Δ::kanMX ctf1Δ::ura4$^+$* | This study |
| AS2197 | *asp1Δ::kanMX dis2Δ::ura4$^+$* | This study |
| AS2198 | *asp1Δ::kanMX rhn1Δ::natMX* | This study |
| AS2207 | *asp1-D333A::kanMX ctf1Δ::ura4$^+$* | This study |
| AS2208 | *asp1-D333A::kanMX dis2Δ::ura4$^+$* | This study |
| AS2217 | *asp1-D333A::kanMX rhn1Δ::natMX* | This study |
| AS2284 | *asp1-H397A::kanMX aps1Δ::natMX ppn1Δ::hygMX* | This study |
| AS2285 | *asp1-H397A::kanMX aps1Δ::natMX swd22Δ::hygMX* | This study |
| AS2309 | *asp1-H397A::kanMX aps1Δ::hygMX ctf1Δ::ura4$^+$* | This study |

| | | |
|---|---|---|
| AS2308 | *asp1-H397A::kanMX aps1Δ::hygMX ssu72-C13S::natMX* | This study |
| AS2304 | *asp1-H397A::kanMX aps1Δ::hygMX rpb1-T4A*$_{29}$*::natMX* | This study |
| AS2337 | *asp1-H397A::kanMX ssu72-C13S::natMX dis2Δ::ura4*$^{+}$ | This study |
| AS2204 | *asp1-D333A::kanMX rpb1-S7A*$_{29}$*::natMX* | This study |
| AS2202 | *asp1-H397A::kanMX rpb1-T4A*$_{29}$*::natMX* | This study |
| AS2290 | *rpb1-S7A*$_{29}$*::natMX ppn1Δ::hygMX aps1Δ::kanMX* | This study |
| AS2291 | *rpb1-S7A*$_{29}$*::natMX swd22Δ::hygMX aps1Δ::kanMX* | This study |
| AS2292 | *rpb1-S7A*$_{29}$*::natMX rhn1Δ::hygMX aps1Δ::kanMX* | This study |
| AS2323 | *rpb1-S7A*$_{29}$*::natMX dis2Δ::ura4*$^{+}$ *aps1Δ::kanMX* | This study |
| AS2324 | *rpb1-S7A*$_{29}$*::natMX ctf1Δ::kanMX aps1Δ::hygMX* | This study |
| AS2325 | *rpb1-S7A*$_{29}$*::natMX ssu72-C13S::kanMX aps1Δ::hygMX* | This study |
| AS2293 | *asp1-H397A::kanMX rpb1-S7A*$_{29}$*::natMX ppn1::hygMX* | This study |
| AS2294 | *asp1-H397A::kanMX rpb1-S7A*$_{29}$*::natMX swd22::hygMX* | This study |
| AS2295 | *asp1-H397A::kanMX rpb1-S7A*$_{29}$*::natMX rhn1Δ::hygMX* | This study |
| AS2313 | *asp1-H397A::kanMX rpb1-S7A*$_{29}$*::natMX ctf1Δ::ura4*$^{+}$ | This study |
| AS2314 | *asp1-H397A::kanMX rpb1-S7A*$_{29}$*::natMX dis2Δ::ura4*$^{+}$ | This study |
| AS2328 | *asp1-H397A::kanMX rpb1-S7A*$_{29}$*::natMX ssu72-C13S::hygMX* | This study |

Figure S7. ***S. pombe* strains used in this study**.
